# Neurocomputational mechanisms underlying immoral decisions benefiting self or others

**DOI:** 10.1101/832659

**Authors:** Chen Qu, Yang Hu, Zixuan Tang, Edmund Derrington, Jean-Claude Dreher

## Abstract

Immoral behavior often consists of weighing transgression of a moral norm against maximizing personal profits. One important question is to understand why immoral behaviors vary based on who receives specific benefits and what are the neurocomputational mechanisms underlying such moral flexibility. Here, we used model-based fMRI to investigate how immoral behaviors change when benefiting oneself or someone else. Participants were presented with offers requiring a tradeoff between a moral cost (i.e., profiting a morally bad cause) and a benefit for either oneself or a charity. Participants were more willing to obtain ill-gotten profits for themselves than for a charity, driven by a devaluation of the moral cost when deciding for their own interests. The subjective value of an immoral offer, computed as a linear summation of the weighed monetary gain and moral cost, recruited the ventromedial prefrontal cortex regardless of beneficiaries. Moreover, paralleling the behavioral findings, this region enhanced its functional coupling with mentalizing-related regions while deciding whether to gain morally-tainted profits for oneself vs. charity. Finally, individual differences in moral preference differentially modulated choice-specific signals in the dorsolateral prefrontal cortex according to who benefited from the decisions. These findings provide insights for understanding the neurobiological basis of moral flexibility.

## 1. Introduction

In almost all cultures and societies human beings tend to transgress established moral values to obtain material advantages in favor of oneself (Bazerman and Gino, 2012; Gächter and Schulz, 2016; Cohn *et al*., 2019). This immoral behavior often consists of weighing motives to uphold a moral norm (e.g., honesty, fairness) against the maximization of personal profits. However, our moral standards change in different contexts. There are numerous examples that demonstrate the remarkable malleability of individuals’ immorality. It is intriguing that the decision to engage in immoral actions varies depending on whether the action benefits oneself or someone else. For instance, people lie more readily when the lie benefits a charity than when it benefits themselves (Lewis *et al*., 2012). In contrast, the magnitude of dishonesty has been observed to increases over time when it benefits oneself but not when it harms oneself while benefitting others (Garrett *et al*., 2016). People also tend to judge others’ moral transgressions (e.g., unfairness) more harshly than their own (Valdesolo and DeSteno, 2007, 2008). Although recent model-based neuroimaging studies have greatly improved our understanding of the neural substrates of (im)morality per se (Hutcherson *et al*., 2015; Crockett *et al*., 2017), the neurocomputational mechanisms that guide flexible immoral decision making depending on the beneficiaries of immoral actions remains poorly understood.

Why do people vary their immoral behaviors depending on who receives the benefits, even when, as perpetrators, they receive no punishment for their behavior? According to the self-concept maintenance theory (Mazar *et al*., 2008), people are often torn between two competing motivations: gaining from immoral actions *versus* maintaining their positive self-concept as a moral individual (Aronson, 1969; Baumeister, 1998; Mazar *et al*., 2008). Thus, individuals may perform immoral actions to benefit themselves financially at the expense of moral self-concept, or, they may forgo financial benefits to maintain their moral self-concept. In order to resolve this moral dilemma, the theory proposes that people often incorporate a level of immorality into their behavior that can be described as “just enough”. This strategy allows for a balance between maintaining a relatively intact self-concept (i.e., I am a morally good person) and pursuing personal profit (Mazar *et al*., 2008; Shalvi *et al*., 2011). This behavioral theory provides a useful framework for investigating flexible immorality. It explains the reason that people vary their moral standards, i.e., the relative weight of financial gain and moral cost differ depending on who benefits from the immoral actions. However, a computational account of this theory that specifies how people vary the trade-offs between monetary gains and moral costs according to the characteristics of the beneficiaries has yet to be defined. Here, we developed and compared computational models of moral decisions incorporating the beneficiary (self/other) of an immoral action, elucidating which variables are computed how they interact and how they are implemented in the brain during immoral decision-making.

At the brain system level, a substantial body of literature from social neuroscience and value-based decision-making established a consensus that the ventromedial prefrontal cortex (vmPFC) plays a key role in value computation for different types of goods (Padoa-Schioppa, 2011; Sescousse *et al*., 2013). Exactly how this region is involved in value computation concerning decisions in social contexts is still debated. While the vmPFC may construct subjective values during decision-making across domains, irrespective of contexts (Levy and Glimcher, 2012; Bartra *et al*., 2013; Ruff and Fehr, 2014), other evidence has revealed its unique involvement when people decide for themselves rather than their partners in a delay-discounting task (Nicolle *et al*., 2012). It is therefore important to directly test whether the vmPFC is engaged in the same way independently of whether it is self or another that benefits from one’s immoral action. In addition to the mass-univariate approach to identify common (or different) neural correlates of the value computation in (or between) the two conditions, we also adopted a representational similarity analysis approach (RSA; Kriegeskorte *et al*., 2008) to assess whether the neural patterns of vmPFC during value computation are similar in the two contexts. The RSA takes advantage of information of multiple voxels to describe the neural pattern similarity between conditions (Kriegeskorte *et al*., 2008; Kriegeskorte and Kievit, 2013), and this approach has been recently applied to the field of social and decision neuroscience (van Baar *et al*., 2019).

Here, we developed a novel paradigm in which participants were asked to make a series of decisions involving trade-offs (i.e., offers) between two parties in the MRI scanner. One party had been established to be a morally bad cause in the opinions of the participants, namely an organization severely violating the moral values of caring for the safety and life of others (a gun-holding/hunting advocacy group; see **SI: association selection** for details). The other party (i.e., the beneficiary) was either the participant (self) or a charity considered to be morally positive. Crucially, by accepting offers in both types of dilemma, either the participant or the charity would be better off; however, it was always accompanied by the moral cost of also profiting the bad cause. When offers were rejected, neither party earned any benefit offered (**Fig. 1**). Notably, to capture how individuals weighs the financial gain and moral cost depending on beneficiaries, we independently varied the monetary payoff for each party (i.e. self/charity vs bad cause) in a parametric manner.

**Fig. 1.**
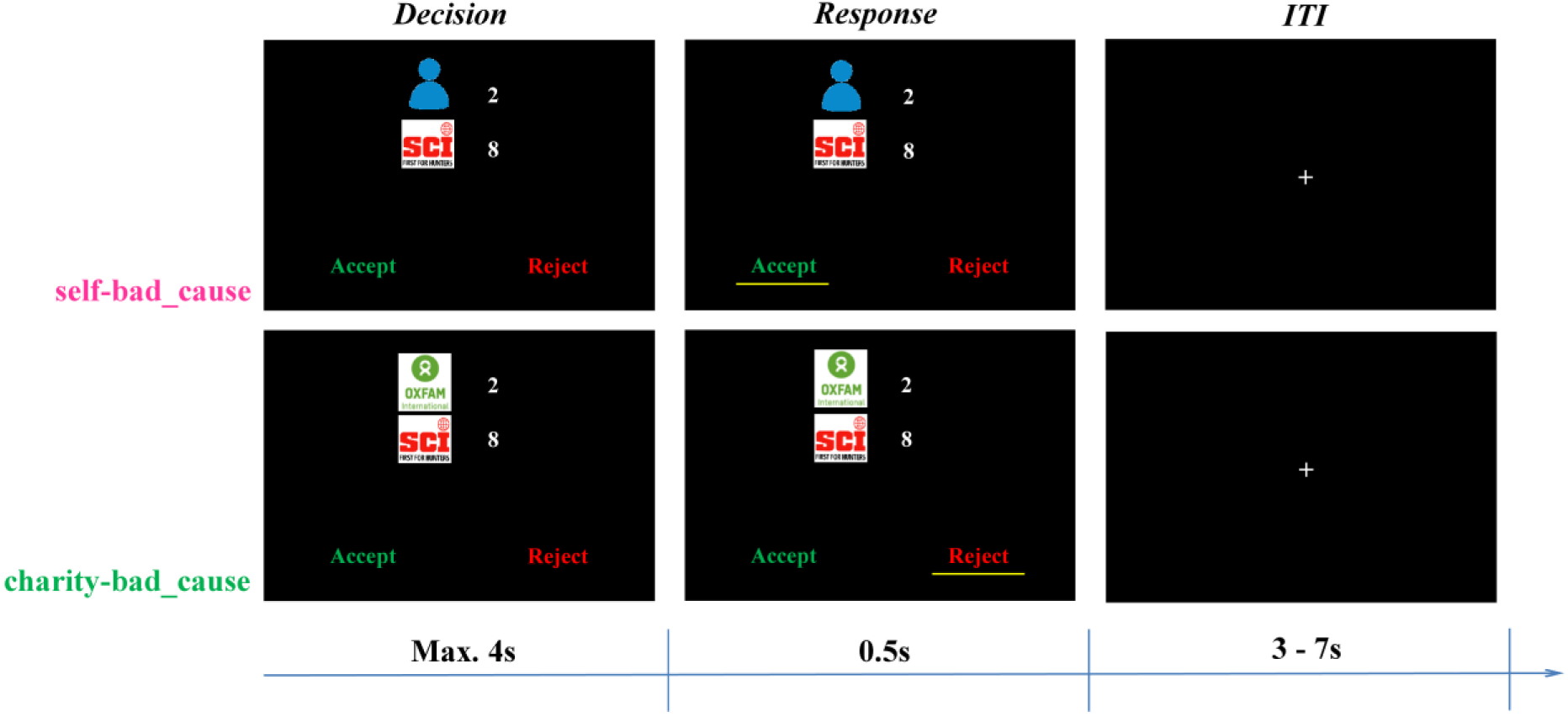
Experimental design. In each trial, participants were first presented with an immoral offer, in which monetary payoffs for two parties were orthogonally varied. One of the parties was always a morally bad cause (i.e., Safari Club International, SCI), whereas the other party was either the participant himself (i.e., a *self-bad cause* dilemma) or the preferred charity (i.e., Oxford Committee for Famine Relief, OXFAM; a *charity-bad cause* dilemma). Participants needed to decide whether to accept or reject the offer within 4 s. If they accepted the offer, both themselves/the charity and the morally bad cause would earn the money as proposed. Otherwise neither party would profit. Each trial was ended with an inter-trial interval (ITI) showing a jittered fixation (3 ∼ 7 s).

This novel experimental design allows us to go beyond traditional behavioral analyses by proposing a series of computational models to elucidate how the human brain computes a decision value integrating moral values and monetary payoffs, and to provide a mechanistic account of flexible immoral choices. We tested and compared a number of such models assuming that immoral choices are made by computing an overall subjective value as a weighted combination of monetary gains for oneself or the charity and the moral cost of benefiting the morally bad cause. This type of value calculation captures a wide range of behavioral patterns in moral choice (Crockett *et al*., 2014; Zhu *et al*., 2014; Volz *et al*., 2017). By comparing the weights related to different beneficiaries, we were able to characterize the computational processes underlying flexible immoral choices.

Moreover, it is of key importance to establish how the vmPFC, likely to compute subjective value of an immoral action, interacts with other brain regions during immoral decisions and whether such functional connectivity changes dependent on the beneficiary of the immoral decision. Previous neuroimaging studies have shown increased functional connectivity between the vmPFC and the temporoparietal junction (TPJ) in a charity-donation task (Hare *et al*., 2010) and in a self-other money-split task (Strombach *et al*., 2015), which consisted of a trade-off between self-profit and benefiting others. We thus investigated whether the functional connectivity between the vmPFC and TPJ is increased when the decider’s own interest is not involved at all in an immoral context.

Additionally, this model-based fMRI approach enabled us to understand the links between inter-subject variability regarding flexible immorality and brain activity depending on the beneficiaries of the morally bad action. More precisely, we aimed to use the model parameters to characterize moral preference across participants (i.e., some subjects are more altruistic and others more selfish), and then to investigate how such inter-individual difference influences the brain activity when accepting that oneself or a charity would benefit (monetarily) from profiting a bad cause (moral cost). It is clear from previous studies that immoral behavior varies from person to person: some subjects never cheat or always cheat but most only cheat sometimes (Fischbacher and Föllmi-Heusi, 2013; Rosenbaum *et al*., 2014). At the brain level, a causal role of the lateral prefrontal cortex (lPFC) has been reported to be the representation of moral goal pursuit (Carlson and Crockett, 2018). Enhancing activity of the right dorsal lPFC (dlPFC) via anodal transcranial direct current stimulation (tDCS) significantly reduced the probability of cheating, providing a causal role of the lPFC in gating immoral behaviors (Maréchal *et al*., 2017). Furthermore, neural signals in lPFC often predict inter-individual difference in immoral behaviors such as self-serving lying (Dogan *et al*., 2016; Yin and Weber, 2018). In a different setting, Crockett and colleagues (2017) revealed an association between lPFC response to immoral earning and inter-individual differences in other-oriented harm aversion. In the light of this literature, we hypothesized that the correlation between inter-individual differences in moral preference and dlPFC activity observed when accepting moral dilemma would depend upon the beneficiary of the immoral choice.

## 2. Methods

### 2.1 Participants

Forty undergraduate or graduate students (25 females; mean age: 20.0 ± 2.0 years, ranging from 18 to 27 years; 2 left handed) were recruited *via* online fliers for the fMRI experiment. All participants had normal or corrected-to-normal vision and reported no prior history of psychiatric or neurological disorders. The study took place at the Imaging Center of South China Normal University and was approved by the local ethics committee. All experimental protocols and procedures were conducted in accordance with the IRB guidelines for experimental testing and were in compliance with the latest revision of the Declaration of Helsinki (BMJ 1991; 302: 1194).

### 2.2 Stimuli

Four charities (i.e., *Oxford Committee for Famine Relief*, *Save the Children*, *First Aid Africa* and *Oceania*) and four non-profit associations advocating gun rights/hunting (i.e., *American Rifle Association*, *The Society for Liberal Weapons Rights*, *Safari Club International*, *The European Federation of Associations for Hunting and Conservation of the EU*) were selected as the charities and morally bad causes for the current fMRI study respectively, based on the ratings by an independent group of participants (N = 30; see **SI: association selection** for details).

The payoff matrix used in the current fMRI study consisted of 64 different combinations between the monetary gain for participants themselves or their preferred charity (i.e., 1 to 8 in steps of 1 monetary unit; 1 MU = CNY 9, same below) and the moral cost (i.e., payoffs for the pre-selected morally bad cause: 4 to 32 in step of 4 MU). The ratio between the monetary gain and the moral cost was deliberately set to 1:4 given previous studies in our lab (Obeso *et al*., 2018) as well as the results of the pilot study (see **SI: pilot behavioral study** for details).

### 2.3 Task

Before scanning, participants were asked to choose one charity and one morally bad association respectively among the 4 candidates mentioned above. Similarly, they read the vignette (with logo) first and then indicated the degree of familiarity (0 = “not at all”, 10 = “very much”) as well as likeness (−10 = “not at all” or “very negative”, 0 = “no preference”, 10 = “very much” or “very positive”) towards each association on a Likert rating scale. To rule out the possibility of equal preference, we also explicitly asked them to indicate the charity (bad cause) he/she likes (dislikes) the most and feels the most (least) willingness to donate.

An event-related design was adopted in the present fMRI study, including 128 trials in total with half in each of two conditions (see below). Trials were presented pseudo-randomly by using M-sequence to improve the efficiency of estimation of hemodynamic responses (Buračas and Boynton, 2002). On each trial, participants were presented with an immoral offer, benefiting two parties with different amount of monetary payoffs. The selected morally bad cause always earned the money, whereas the other party was either participants themselves (i.e., a *self-bad cause* dilemma) or the preferred charity (i.e., a *charity-bad cause* dilemma). In either dilemma, participants were faced up with two options, i.e., “accept” or “reject”, with the position counterbalanced across participants but fixed within the participant. If they chose to accept the offer, both themselves/the charity and the morally bad cause would earn the money as proposed. Otherwise neither parties would profit. Participants were asked to respond within 4 s by pressing a corresponding button on the button box with the left/right index finger. If an invalid response was made (i.e., no response in 4 s or response less than 200 ms), a warning screen showed up and this trial was repeated at the end of the scanning session. Each trial ended up with an inter-trial interval showing a jittered fixation (3 ∼ 7 s).

Participants were told that their decisions were independent from trial to trial and that once the present task was chosen to be paid (see procedure for details), one trial in each dilemma would be randomly selected to determine their final payoff and the corresponding donation made to the preferred charity. The final amount donated to the pre-selected morally bad cause was randomly determined between one of the two selected trials mentioned above. In fact, we only paid participants accordingly and no donations were made to these associations. Participants were informed of this at the very end of the experiment.

### 2.4 Procedure

On the day of scanning, participants signed a written informed consent and were explained the procedure, which included the present task and another independent task which will be reported elsewhere To rule out the possibility of hedging the income risk across two tasks, they were informed that besides the participation fee (i.e., 80 CNY ≈ 12.7 USD), only one task, randomly chosen by the computer at the end of the experiment, would be paid in addition to their basic fee.

For the current task, participants were first provided with the instructions and then they selected their favorite charity as well as the morally bad cause they disliked the most. Before the fMRI task, participants completed a series of comprehension questions to ensure that they fully understood the task and also performed a practice session to get familiar with the paradigm as well as the response button in the scanner. The scanning part included one functional session lasting around 15 min, which was followed by a 6-min structural scan. After that, participants filled out a battery of questionnaires by indicating degrees on a Likert rating scale, including the degree of moral conflict when they made the decisions (0 = “not at all”, 100 = “very much”) and that of moral inappropriateness if they accepted the offer (0 = “not at all”, 100 = “very much”), for each dilemma separately. They also filled out several scales of personality traits used for the exploratory analyses. After completing this, participants were debriefed, paid, and acknowledged.

### 2.5 Behavioral Analyses

All behavioral analyses were conducted using R (http://www.r-project.org/) and relevant packages (R Core Team, 2014). All reported p values are two-tailed and p < 0.05 was considered statistically significant. Data visualization were performed via “ggplot2” package (Wickham, 2016).

Regarding the choice data, we performed a repeated mixed-effect logistic regression on the decision of choosing the “accept” option by the *glmer* function in “lme4” package (Bates *et al*., 2013), with dilemma (dummy variable; reference level: *self-bad cause* dilemma; same below) and payoffs for both parties involved in each dilemma (i.e., the monetary gain and the moral cost; mean-centered continuous variable; same below) as the fixed-effect predictors. In addition, we included the following random-effect factors allowing varying intercept across participants. For the statistical inference on each predictor, we performed the Type II Wald chi-square test on the model fits by using the *Anova* function in “car” package (Fox *et al*., 2016), and reported the odds ratio as relevant effect size.

For decision time (DT), we first did a log-transformation due to its non-normal distribution (Anderson-Darling normality test: A = 91.90, p < 0.001) and then performed a mixed-effect linear regression on the log-transformed DT by the *lmer* function in “lme4” package, with decision (dummy variable; reference level: accept), dilemma, decision × dilemma, as well as payoffs for both beneficiaries as the fixed-effect predictors. Random-effect factors were specified in the same way as above. Similar analyses were also performed on the post-scanning rating except that dilemma was added as the only fixed-effect predictor. We followed the procedure recommended by Luke (2017) to obtain the statistics for each predictor by applying the Satterthwaite approximations on the restricted maximum likelihood model (REML) fit via the “lmerTest” package (Luke, 2017). In addition, we computed the Cohen’d of each predictor via the “EMAtools” package (Kleiman, 2017), which provided the effect size measure specially for the mixed-effect regressions. For likeness ratings of the selected associations, we compared whether the ratings significantly differed from 0 in each type of selected associations (i.e., charity or morally bad causes) respectively by the one-sample T-test, and computed the Cohen’s d as effect size.

### 2.6 Computational Modelling

To examine how participants integrated the payoffs of both parties in two different dilemmas into a subjective value (SV), we compared the following 10 models with different utility functions.

Model 1 was adapted from a recent study on moral decision making by Crockett *et al* (2014, 2015, 2017). The model described that the subjective value was calculated by the gains for participants or the charity relative to that for the bad cause, which could be formally represented as follows:

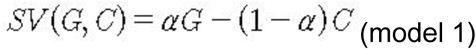

where SV denotes the SV of the given trial if the participant chooses to accept. For rejection trials, SV is always 0 given the rule of the task (i.e., neither beneficiaries would gain the money regardless of dilemmas; same for all models). **G** represents the monetary **G**ain for the participant or the charity, while **C** represents the moral cost, measured by the monetary gain to the morally bad cause. α is the unknown parameter of social preference that characterizes the relative weight on the payoff for either party involved in the dilemma (0 < α < 1).

Model 3 was based on the study by Park *et al* (Park *et al*., 2011) which initially examined the integration of positive and negative values and recently was adapted to a donation task (Lopez-Persem *et al*., 2017):

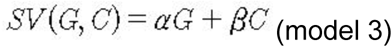

where α and β are unknown parameters which capture the weight of the payoff for either party involved in the dilemma (−10 < α < 10, −10 < β < 10).

Model 5 was based on the Fehr-Schmidt model (Fehr and Schmidt, 1999):

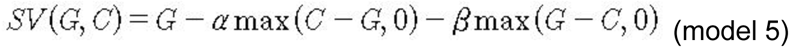

where α and β measure the degree of aversion to payoff inequality in disadvantageous and advantageous situation respectively (i.e., how much participants dislike when they themselves/the charity earned less/more than the bad cause; 0 < α < 5, 0 < β < 1).

In addition, we also included Model 7 assuming that people are aversive to the absolute payoff inequality between two beneficiaries, captured by a parameter θ (0 < θ < 5):

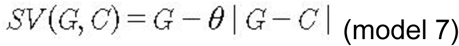

Models with even index mimic corresponding models with odd index (i.e., Model 2, 4, 6, 8 matches with Model 1, 3, 5, 7 respectively) except that those unknown parameters varied dependently on the two dilemmas.

The probability of choosing the reject option was determined by the softmax function:

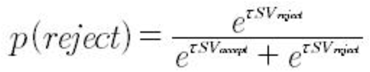

where τ refers to the inverse softmax temperature (0 < τ < 5) which denotes the sensitivity of individual’s choice to the difference in SV between options of acceptance and rejection.

We used the “hBayesDM” package (Ahn *et al*., 2017) to fit all aforementioned candidate models using the Hierarchical Bayesian Analysis (HBA) approach (Gelman *et al*., 2014). The “hBayesDM” package is developed based on the Stan language (Stan Development Team, 2016) which utilizes a Markov Chain Monte Carlo (MCMC) sampling scheme to perform full Bayesian inference and obtain the actual posterior distribution. We adopted HBA rather than maximum likelihood estimation (MLE) because HBA provides more stable and accurate estimates than MLE (Ahn *et al*., 2011). Following the approach in “hBayesDM” package, we assumed the individual-level parameters ϕ were drawn from a group-level normal distribution: *ϕ* ∼ *Normal* (*μ_ϕ_*, *σ_ϕ_*), where *μ_ϕ_* and *σ_ϕ_* refer to the group-level mean and standard deviation respectively. Weakly-informative priors were adopted for both these group-level parameters, i.e., *μ_ϕ_* ∼ *Normal* (0, 1) and *σ_ϕ_* ∼ *half-Cauchy* (0, 2) (Ahn *et al*., 2017). In HBA, all group-level parameters and individual-level parameters are simultaneously estimated through the Bayes rule given the behavioral data. We fit each candidate model with four independent MCMC chains using 1,000 iterations after 1,500 iterations for initial algorithm warmup per chain, resulting in 4,000 valid posterior samples. Convergence of the MCMC chains was assessed through the Gelman-Rubin R-hat Statistics (Gelman and Rubin, 1992).

For model comparison, we computed the widely applicable information criterion (WAIC) score per candidate model (Vehtari *et al*., 2016). WAIC score provides the estimate of out-of-sample predictive accuracy in a fully Bayesian way, which is more reliable compared to the point-estimate information criterion (e.g., AIC). By convention, the lower WAIC score indicates better out-of-sample prediction accuracy of the candidate model. A difference score of 10 on the information criterion scale is considered decisive (Burnham and Anderson, 2004). We selected the model with the lowest WAIC as the winning model for subsequent analysis. In addition, we also implemented a posterior predictive check to further examine the absolute performance of the winning model, i.e., whether the prediction of the winning model could characterize the feature of real choices. In specific, we employed each individual’s joint posterior MCMC samples (i.e., 4,000 times) to generate new choice datasets correspondingly (i.e., 4,000 choices per trial per participant), given the actual trial-wise stimuli sequences presented to each participant. Thus we obtained the model prediction by calculating the average rejection proportion of these new datasets in terms of two dilemmas for each subject respectively. We tested to what degree the individual model prediction correlated with the actual rejection proportion using Pearson correlation. Based on the winning model and its parameter estimation, we derived the mean of the trial-wise subjective value (SV) for each option and defined the relative SV (rSV) by subtracting the SV of the non-chosen option from that of the chosen option (i.e., rSV = SV_chosen - SV_unchosen). These trial-wise rSV were used as parametric modulators for model-based fMRI analyses (see below for details).

### 2.7 fMRI Data Acquisition and Analyses

The imaging data were acquired on a 3-Tesla Siemens Trio MRI system (Siemens, *Erlangen, Germany*) with a 32-channel head coil at the Imaging Center of South China Normal University. Functional data were acquired using T2*-weighted echo-planar imaging (EPI) sequences employing a BOLD contrast (TR = 2000 ms, TE = 30 ms; flip angle = 90°; slice thickness = 3.5 mm, slice gap = 25%, matrix = 64 × 64, FoV = 224 × 224 mm^2^) in 32 axial slices. Slices were axially oriented along the AC-PC plane and acquired in an ascending order. A high-resolution structural T1-weighted image was also collected for every participant using a 3D MRI sequence (TR = 1900 ms, TE = 2.52 ms; flip angle = 9°; slice thickness = 1 mm, matrix = 256 × 256, FoV = 256 × 256 mm²).

Three participant were excluded from later analyses due to excessive head movements (> 3 mm), thus leaving a total of 37 participants whose data were analyzed for the fMRI study (24 females; mean age ± SD = 19.9 ± 2.0 years, ranging from 18 to 27 years; 2 left handedness). Functional imaging data were analyzed using SPM12 (Wellcome Trust Centre for Neuroimaging, University College London, *London, UK*). The preprocessing procedure followed the pipeline recommended by SPM12. In particular, functional images (EPI) were first realigned to the first volume to correct motion artifacts, unwarped, and corrected for slice timing. Next, the structural T_1_ image was segmented into white-matter, grey-matter and cerebrospinal fluid with the skull removed, and co-registered to the mean functional images. Then all functional images were normalized to the MNI space, resampled with a 2 × 2 × 2 mm^3^ resolution, based on parameters generated in the previous step. Last, the normalized functional images were smoothed using an 8-mm isotropic full width half maximum (FWHM) based on Gaussian kernel.

#### 2.7.1 General Linear Models (GLMs) Analyses

For all GLMs below, the canonical hemodynamic response function (HRF) was used and a high-pass temporal filtering was performed with a default cut-off value of 128 s to remove low-frequency drifts.

For each participant, we constructed the following GLMs. GLM1 focused on investigating brain regions encoding the relative subjective value (rSV) which integrates the monetary gain and moral cost during decision-making period in each dilemma. Thus, we included the following regressors of interest, namely onsets of the decision period in each dilemma with the duration of actual decision time (DT). Each regressor of onset was associated with the rSV based on the winning model as the parametric modulators (PMs). For the completeness of analyses, we also established GLM2 to identify regions parametrically encoding monetary gain and moral cost during the decision period in each dilemma. We included the same regressor of onsets as in GLM1, except that each regressor of onsets associated with two PMs, i.e., the monetary gain (*self-bad cause* dilemma: payoff for the participant; *charity-bad cause* dilemma: payoff for the charity) and the moral cost (in both dilemmas: payoff for the bad cause). Notably, the default orthogonalization option in SPM12 was switched off to ensure the competition for variance during estimation of two PMs. For these two GLMs, we built up the contrasts of each PM against implicit baseline, and that between two dilemmas for the group-level analyses. GLM3 was established to estimate the choice-specific neural activities in different dilemmas (i.e., the dilemma × decision interactive signals). Four participants were excluded from this analyses due to the missing accept (N = 2) or reject (N = 2) decisions in the *charity-bad cause* dilemma. GLM3 was constructed in the same way as GLM2, except that we sorted the onsets of decision period by different decision in each dilemma. We built up the dilemma × decision contrasts (i.e., differential neural activities between accept vs. reject between two dilemmas) for the group-level analyses.

Regarding the regressors of non-interest, we modelled the onset of button press to rule out the movement effect for all GLMs. Besides, once the participant showed invalid responses, an additional regressor modelling relevant events (i.e., *other*) was included, which contained decision onsets of invalid trials (i.e., for trials which DT are less than 200 ms, duration equals the actual DT; for trials of no response, duration equals 4s) as well as the warning feedback (duration equals 1s). Furthermore, the six movement parameters were added to all models as covariates to account for artifact of head motion.

#### 2.7.2 Representational Similarity Analyses (RSA)

The RSA was carried out in Python 3.6.3 with the NLTools package (v.0.3.6; http://github.com/ljchang/nltools), which aimed to further examine whether the neural patterns in vmPFC during value computation in the *self-bad cause* dilemma could mimic the one in the *charity-bad cause* dilemma. For each participant, we established a neural dissimilarity matrix (DM) within the vmPFC (defined based on the conjunction activation in two dilemmas; see Results for details) between value-related contrast maps in the *self-bad cause* dilemma and in the *charity-bad cause* dilemma (i.e., the PM contrasts of rSV in GLM 1 in respective dilemmas). The neural DM was calculated by 1 minus the Pearson correlation between contrast value vectors of vmPFC pattern of two dilemmas. Next, we transformed the individual dissimilarity value back to the correlation coefficient, and then performed the Fisher z transformation for statistical analyses. Besides one-sample t-test, we also did permutation analysis by shuffling individual labels and running the same analysis for 5000 times, and finally calculated the proportion of cases as the significance level of such correlation in which the permuted mean z-value exceeded the true mean z-value.

#### 2.7.3 Functional Connectivity Analyses

To address how the functional connectivity between the region encoding value-signals (i.e., vmPFC) and the rest of brain changes between dilemmas, by taking a generalized psychophysical interaction (gPPI) approach (McLaren *et al*., 2012). To this end, for each participant, we constructed a PPI-GLM (based on GLM1) using the gPPI toolbox (https://www.nitrc.org/projects/gppi) 1) to extract the de-convolved time series at the group-level peak of the joint activation of vmPFC (i.e., encoding the rSV in both dilemmas; peak MNI: -2/48/-14; see Results for details) within a 6mm-radium sphere as the physiological regressor, 2) to define all regressors (i.e., onsets and PMs) in GLM1 as the psychological regressors, and 3) to multiply the physiological regressor with each psychological regressor as the PPI regressors. These regressors were all convolved with the canonical HRF to model the BOLD signal. In addition, we also incorporated six movement parameters as covariates to account for artifact of head motion. We then built up the individual-level PPI contrasts between two dilemmas and used them for the group-level analyses.

#### 2.7.4 Statistical Inference and Visualization

Individual-level contrasts mentioned above were fed to the group-level random-effect analyses. One-sample T-tests, conjunction analyses (Nichols *et al*., 2005), and regression analyses were performed to detect the differential neural activities, joint activation, and behavioral-brain correlation respectively. For whole-brain analyses, we adopted p < 0.05 at the cluster-level controlling for family-wise error (FWE) rate combining with an uncorrected voxel-level threshold of p < 0.001 as the analyses of the whole-brain threshold (Eklund *et al*., 2016). Based on our hypotheses, we also adopted the following regions of interest (ROI) for specific contrasts by performing a small volume correction (SVC), i.e., the ventromedial prefrontal cortex (vmPFC: 2/46/-8) related with value-computation (Bartra *et al*., 2013), the temporoparietal junction (TPJ; left: -53/-59/20; right: 56/-56/18) related with mentalizing (Schurz *et al*., 2014), and the dorsolateral prefrontal cortex (dlPFC: ±46/36/24) related with moral judgment and decision-making (Greene *et al*., 2001). All these ROIs were defined by 9-mm spheres with corresponding MNI coordinates as the center. Regions were labelled according to the automated anatomical labelling (AAL) template via the xjView toolbox (http://www.alivelearn.net/xjview8/).

To visualize the effect of PMs on neural activities in relevant ROIs (i.e., vmPFC) over time, we followed the procedure used by (Fleming *et al*., 2018). In brief, we first extracted the de-noised time courses within the masks mentioned above from 10 s windows time-locked to the onset of decision. Then we applied a regression with corresponding standardized PMs (i.e., rSV) to the standardized activity of each time point in each dilemma respectively, resulting in a time course of β weights of PMs. In case of illustrating the effect of dilemma on the modulation of PMs, we ran similar regressions except that we pooled the two conditions together and adopted the dilemma, PMs and their interactions as predictors. We used non-parametric permutation tests (1,000 permutations) to assess group-level significance of β weights against 0. Significant effect for individual time points were marked by asterisks if the actual t-statistic fell outside the 2.5th or 97.5th percentiles of the null distribution generated by the permutation test.

## 3. Results

### 3.1 Behavioral Results

Each candidate for charity and morally bad cause were selected by participants at least once (see **Fig. S1a**). None of the selected associations were familiar to them, as indicated by the low average scores for familiarity (i.e., less than 2 on a 0-10 Likert scale; mean ± SD: charity: 1.68 ± 1.80; morally bad cause: 0.46 ± 1.10). However, participants rated the chosen charities positively (mean ± SD (95% confidence interval (CI)): 8.57 ± 1.52 (8.06, 9.07); t(36) = 34.31, p < 0.001, Cohen’s d = 5.64) and had negative evaluations of the chosen gun/hunting rights advocacy group (i.e., morally bad cause; mean ± SD (95% CI): −8.03 ± 2.65 (−8.91, −7.14); t(36) = −18.42, p < 0.0014, Cohen’s d = 3.03, see **Fig. S1b**).

Although participants stated that it felt less morally inappropriate to accept offers to benefit the charity rather than themselves (48.1 ± 32.5 vs. 59.1 ± 30.0; b (95% CI) = −11.03 (−21.13, −0.93), SE = 5.09, t(36) = −2.167, p = 0.037, Cohen’s d = −0.72) and rated comparable levels of moral conflict during the decision-making period for the self and charity conditions (46.1 ± 32.0 vs. 48.5 ± 29.6; b (95% CI) = −2.43 (−12.68, 7.81), SE = 5.16, t(36) = −0.471, p = 0.640, Cohen’s d = −0.16), their behavior did not tell the same story. Specifically, participants were less likely to accept offers in the *charity-bad cause* dilemma vs. *self-bad cause* dilemma (acceptance rate: 39.7 ± 26.5 % vs. 47.3 ± 35.9 %; Odds Ratio = 0.47, b (95% CI) = −0.75 (−0.93, −0.57), SE = 0.09, χ^2^(1) = 65.26, p < 0.001). We also found that higher monetary gain for the participants themselves or the charity (Odds Ratio = 1.91, b (95% CI) = 0.65 (0.60, 0.70), SE = 0.03, χ^2^(1) = 640.36, p < 0.001) made participants more likely to accept offers whereas the moral cost showed the opposite effect (Odds Ratio = 0.87, b (95% CI) = −0.14 (−0.15, −0.13), SE = 0.01, χ^2^(1) = 520.91, p < 0.001; see **Fig. 2**).

**Fig. 2.**
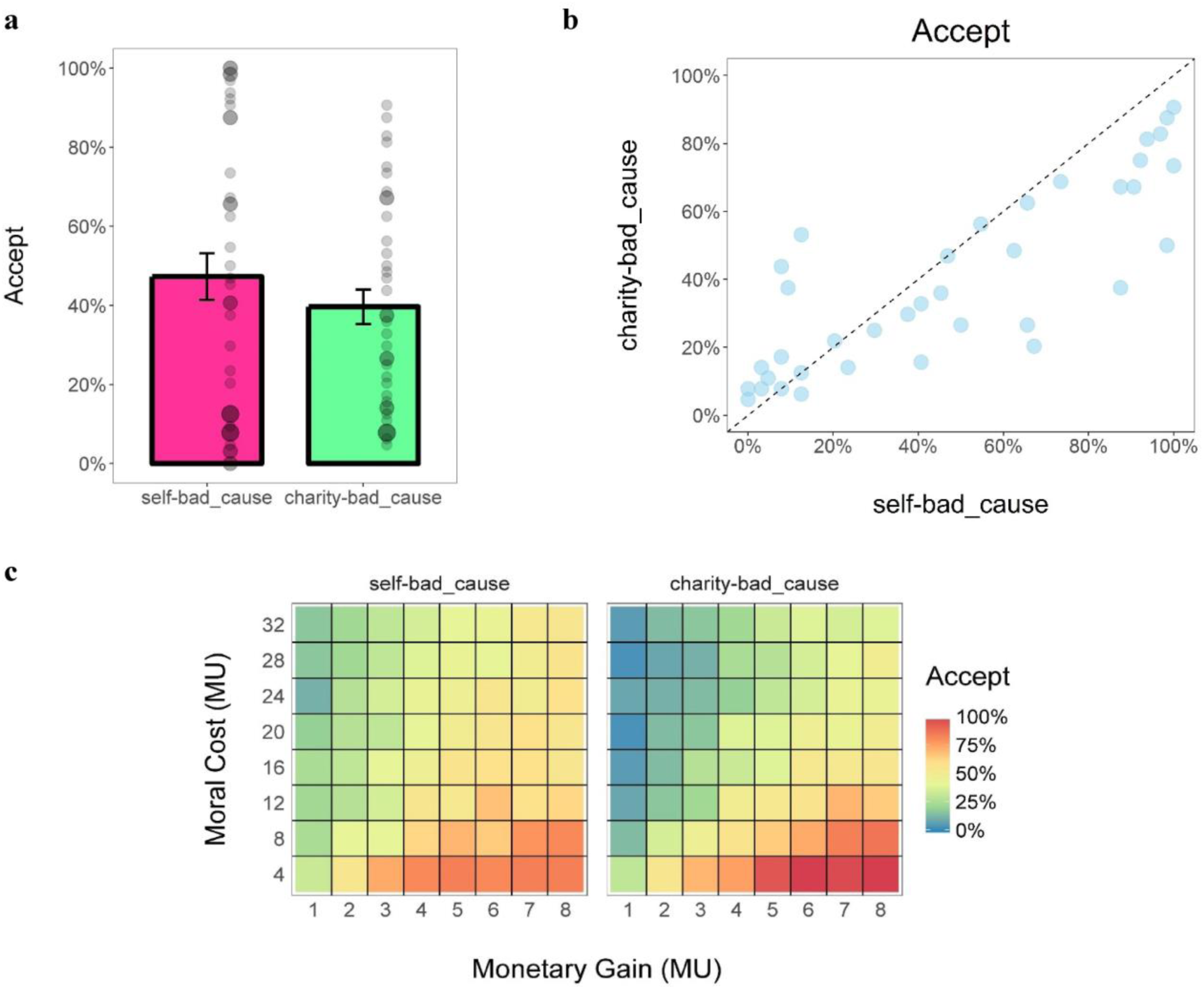
Behavioral results of the fMRI study. (a) Mean acceptance rate in each dilemma. Each dot refers to the acceptance rate of a single participant. (b) Correlation of acceptance rate between two dilemmas. Each dot refers to the acceptance rate of a single participant. Dots below the dotted diagonal indicates participants who accepts more immoral offers in the *self-bad_cause* dilemma than in the *charity-bad_cause* dilemma. (c) Heat map of the mean acceptance rate (%) as a function of the monetary gain and the moral cost in each dilemma. Error bars represent the SEM; Abbreviation: MU = monetary unit.

Concerning the relationship of choice behaviors between the two conditions, we found that participants who accepted more immoral offers in the *self-bad cause* dilemma also accepted offers more frequently in the *charity-bad cause* dilemma (r(95% CI) = 0.830 (0.692, 0.910), t(35) = 8.81, p < 0.001; see **Fig. 2**). Moreover, participants who accepted the previous immoral offer in the *self-bad cause* dilemma were less likely to reject the current immoral offer in the *charity-bad cause* dilemma (Odds Ratio = 0.77, b (95% CI) = −0.26 (−0.45, −0.07), SE = 0.10, χ^2^(1) = 7.08, p = 0.008), after controlling the effect of the monetary gain (Odds Ratio = 0.81, b (95% CI) = −0.21 (−0.24, −0.18), SE = 0.02, χ^2^(1) = 180.69, p < 0.001) and moral cost (Odds Ratio = 1.05, b (95% CI) = 0.05 (0.04, 0.06), SE = 0.004, χ^2^(1) = 159.99, p < 0.001) in the current trial.

For decision time (DT), we first did a log-transformation due to its non-normal distribution (Anderson-Darling normality test: A = 91.90, p < 0.001). Regressions on log-transformed DT revealed a trend-to-significant dilemma × decision interaction (b (95% CI) = −0.04 (−0.07, 0.001), SE = 0.02, t(4702) = −1.87, p = 0.062, Cohen’s d = −0.05; see **Fig. S2**). In addition, higher moral cost accelerated the decision process (b (95% CI) = −0.002 (−0.003, −0.001), SE = 0.0005, t(4079) = −3.05, p = 0.002, Cohen’s d = −0.09). However, participants made decisions more slowly when the earnings for themselves or the charity were large (b (95% CI) = 0.01 (0.008, 0.017), SE = 0.002, t(4714) = 5.54, p < 0.001, Cohen’s d = 0.16). To unpack the marginal significant interaction effect, we ran the same analyses on acceptance and rejection decisions seperately. The effect of dilemma on DT in both acceptance choice (*self-bad cause* dilemma vs. *charity_bad cause* dilemma: 1553.6 ± 473.3 ms vs. 1558.3 ± 366.5 ms; b (95% CI) = 0.09 (0.06, 0.11), SE = 0.01, t(2028) = 6.00, p < 0.001, Cohen’s d = 0.27) and rejection choice (1474.7 ± 304.1 ms vs. 1481.1 ± 278.9 ms; b (95% CI) = 0.02 (0.001, 0.05), SE = 0.01, t(2652) = 2.08, p = 0.038, Cohen’s d = 0.08) after controlling the effect of payoff (acceptance: monetary gain: b (95% CI) = −0.02 (−0.03, −0.01), SE = 0.003, t(2027) = −6.23, p < 0.001, Cohen’s d = −0.28; moral cost: b (95% CI) = 0.005 (0.003, 0.006), SE = 0.0008, t(2039) = 5.84, p < 0.001, Cohen’s d = 0.26; rejection: monetary gain: b (95% CI) = 0.037 (0.031, 0.042), SE = 0.003, t(2659) = 14.06, p < 0.001, Cohen’s d = 0.55; moral cost: b (95% CI) = −0.005 (−0.006, −0.004), SE = 0.0007, t(2643) = −7.64, p < 0.001, Cohen’s d = −0.30).

### 3.2 Computational Modelling Results

We fitted the computational models noted above to the choice data by adopting the Hierarchical Bayesian Analysis (HBA) approach (Gelman *et al*., 2014) via the R package “hBayesDM” (Ahn *et al*., 2017). R-hat values of all estimated parameters of all models were close to 1.0 (at most smaller than 1.03 in the current case), indicating adequate convergence of the MCMC chains (Gelman and Rubin, 1992). The hierarchical Bayesian model comparison showed that the **model 4** below was with lowest WAIC scores and outperformed other competitive models (see **Fig. 3a**):

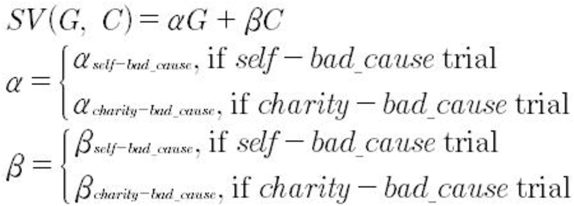

**Fig. 3.**
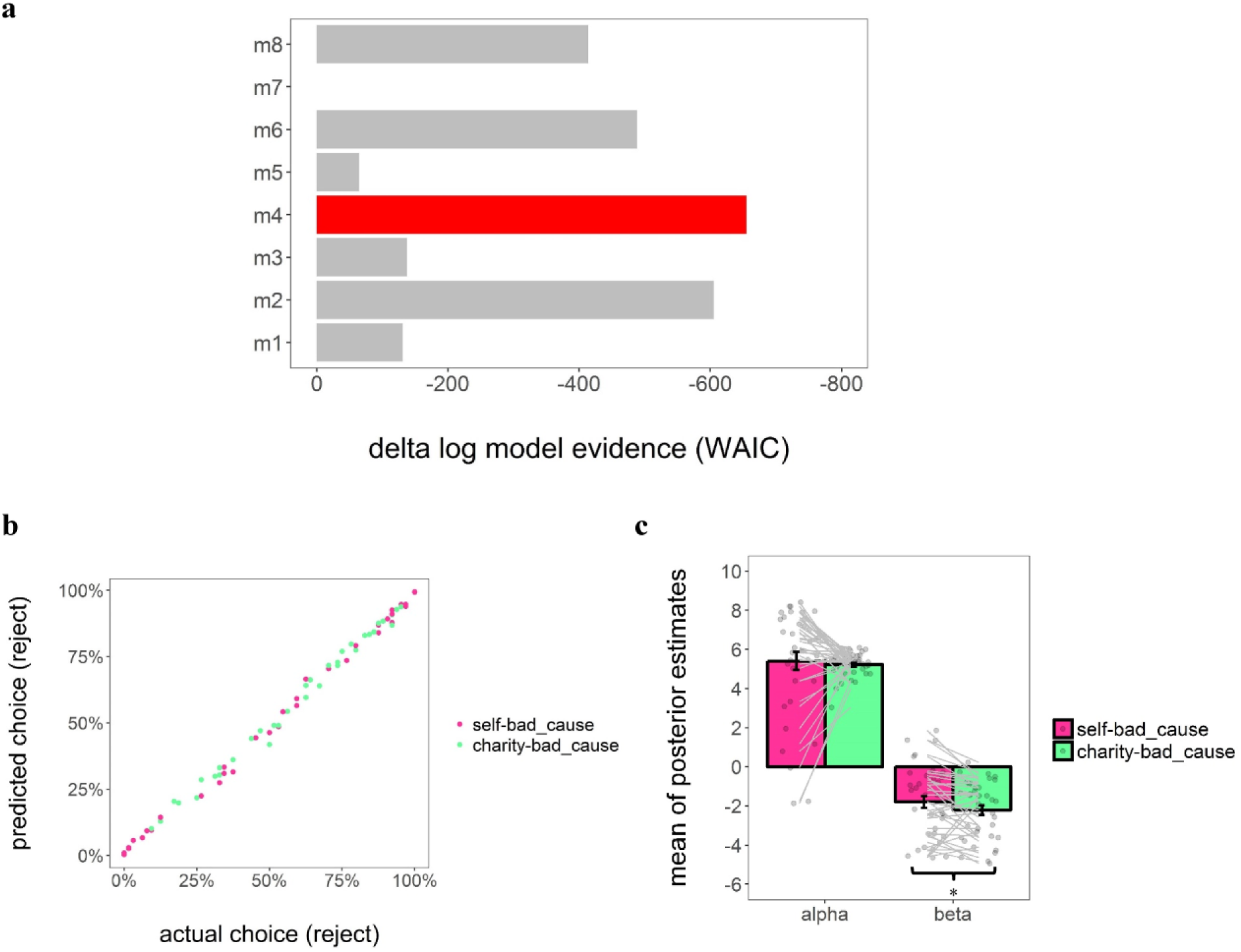
Results of computational modelling. (a) Bayesian model evidence. In our case, model evidence (relative to the model with the worst accuracy of out-of-sample prediction, i.e., model 7) clearly favors to model 4 (m4). Lower (i.e., more negative) Watanabe-Akaike Information Criterion (WAIC) scores indicates better model. (b) Correlation between the actual reject rate (%) and the mean predicted reject rate (%) based on the posterior distribution of the parameters estimated based on the winning model in each dilemma. Each dot represents the data of a single participant. (c) Group-level mean of posterior estimates of individual-level key parameters (i.e., α and β) based on the winning model. Each dot represents the data of a single participant. Error bars represent the SEM; significance: *p< 0.05.

This model was adapted from the study by Park *et al* (Park *et al*., 2011) which initially examined the integration of positive and negative values and recently was adapted to a donation task (Lopez-Persem *et al*., 2017). G and C represent the monetary gain for the participant or the charity and the morally bad cause respectively. α and β are unknown parameters which capture the weight of the monetary gain and moral cost involved in the dilemma respectively (−10 < α < 10, −10 < β < 10). On top of it, this model distinguished weights on payoffs of both parties in terms of dilemma. The posterior predictive check further showed that the prediction of the winning model highly correlated the actual choice behavior (*self-bad cause* dilemma: r(95% CI) = 0.998 (0.996, 0.999), t(35) = 94.06, p < 0.001; *charity-bad cause* dilemma: r(95% CI) = 0.996 (0.993, 0.998), t(35) = 0.996, p < 0.001; see **Fig. 3b**). Taking a closer look at these individual-level posterior mean of key parameters estimated from the winning model, we found that participants weighted positively the monetary gains either for themselves (α_self-bad_cause_: mean ± SD (95% CI): 5.41 ± 2.79 (4.48, 6.34); t(36) = 11.80, p < 0.001, Cohen’d = 1.94) or the charity (α_charity-bad_cause_: mean ± SD (95% CI): 5.22 ± 0.63 (5.01, 5.43); t(36) = 50.41, p < 0.001, Cohen’d = 8.29), whereas they weighted negatively the moral cost in both dilemmas (β_self-bad_cause_: mean ± SD (95% CI): −1.79 ± 1.81 (−2.39, −1.19); t(36) = −6.03, p < 0.001, Cohen’d = 0.99; β_charity-bad_cause_: mean ± SD (95% CI): −2.21 ± 1.47 (−2.70, −1.72); t(36) = −9.12, p < 0.001, Cohen’d = 1.50). Paired-wise t-test further showed that participants weighted the moral cost more negatively in the *charity-bad cause* dilemma vs. *self-bad cause* (95% CI of mean difference: −0.84, −0.01; t(36) = −2.07, p = 0.046, Cohen’s d = 0.34), whereas their weights on the monetary gains were comparable for themselves and the charity (95% CI of mean difference: −1.05, 0.68; t(36) = 0.44, p = 0.662, Cohen’s d = 0.07; see **Fig. 3c**).

To further characterize the inter-individual variance in differential modulatory effect of dilemma on immoral decisions, we computed an index of moral preference in the following way:

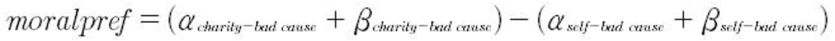

the higher the index is, the stronger the preference of participants to weight monetary gain for the charity higher than for themselves when controlling the weights of the moral cost in the two dilemmas respectively. Notably, we standardized the original payoffs and re-fitted the winning model (i.e., m4) to the dataset. This made the parameter estimates capturing the weights of both the monetary gains and the moral costs comparable on the same scale. As a *post-hoc* check, we also observed a negative correlation between the moral preference and the total score of Machiavelli scale (Mach-IV; Pearson correlation: r (95% CI) = −0.32 (−0.59, −0.001), t(35) = −2.03, p = 0.049; see **Fig. S3**), where a higher score indicates someone who agrees with the views of achieving one’s own purposes or interests by manipulating others even via immoral ways (Christie & Geis, 2013). This finding justified the external validity of the moral preference measured by the present task.

### 3.3 Neuroimaging Results

#### 3.3.1 vmPFC encodes relative subjective values (rSV) during immoral decision-making in both dilemmas (GLM1)

The conjunction analyses of both parametric contrasts (against implicit baseline) showed that the activity in vmPFC was positively modulated by rSV generated based on the winning model (peak MNI coordinates: -2/48/-14, t(72) = 3.08, p(SVC-FWE) = 0.043; see **Fig. 4**; also see **Table S1** for details of other activated regions). This suggests that immoral decisions rely on the same brain circuitry regardless of the beneficiary of the bad action. This was in line with a direct comparison between brain regions modulated by the rSV in the two dilemmas, which did not reveal any significant difference.

**Fig. 4.**
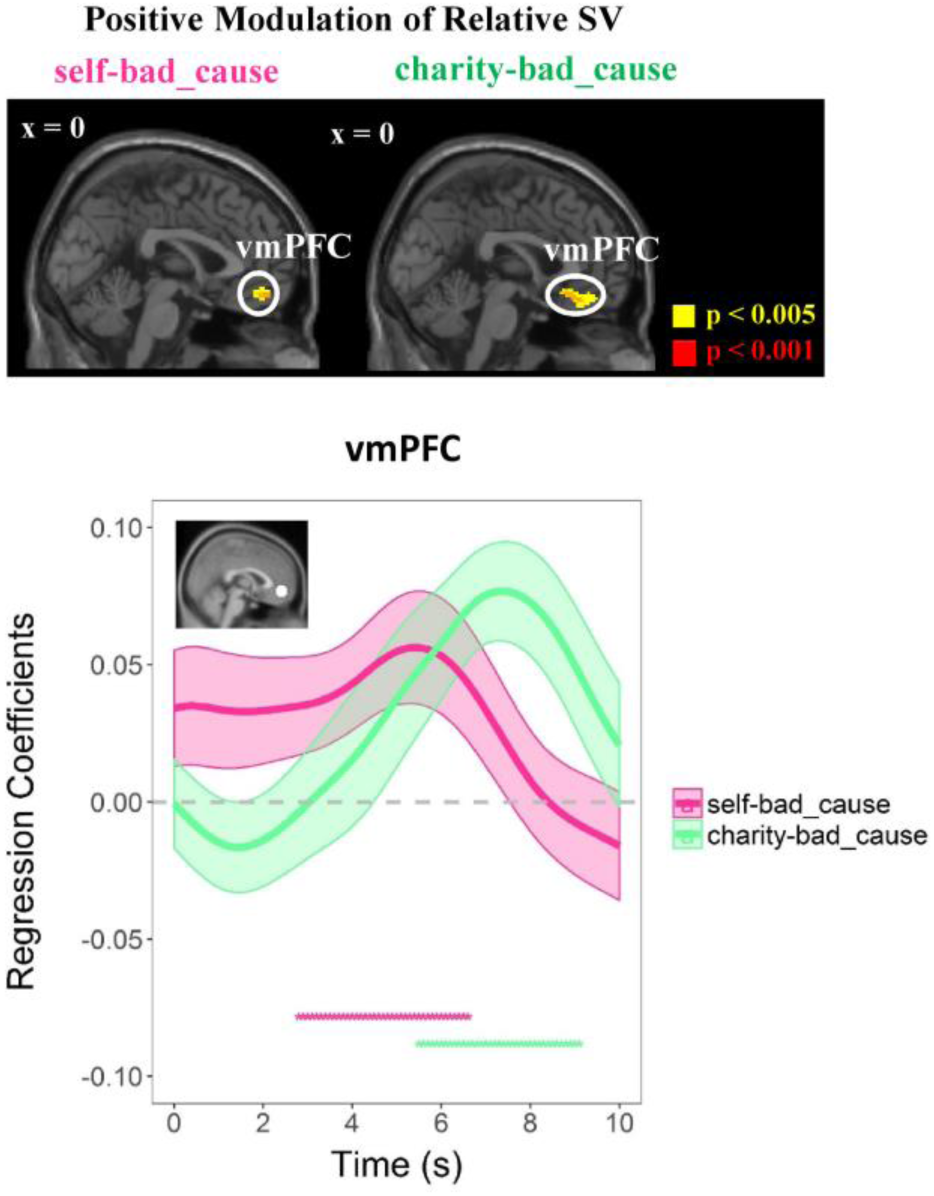
vmPFC encodes relative subjective value (SV) during decision-making period regardless of beneficiaries. Top panel: Positive modulation of relative SV on the ventromedial prefrontal cortex (vmPFC) in both dilemmas (GLM2). Bottom panel: regression analysis of the effects of relative SV on an independent ROI (i.e., Bartra *et al*., 2013: center MNI coordinates: x/y/z = -2/46/-8; a sphere with a radius of 9 mm;) activity time courses in each dilemma; the significant effect was indicated by the magenta or green dots. Regression coefficients are represented as group-level mean ± SEM (shaded areas). Dots below the time course indicate significant excursions of t statistics assessed using two-tailed permutation tests. Display threshold: p < 0.001 and p < 0.005 uncorrected at the voxel-level.

Furthermore, we performed an additional RSA to directly test whether the neural patterns of vmPFC during value computation identified in the *self-bad cause* dilemma is similar to the that in the *charity-bad cause* dilemma. Here, we defined the ROI in vmPFC by constructing a 6 mm sphere with the peak coordinate of the conjunction analysis as the center. Consistent with the conjunction analysis, the neural patterns of vmPFC in two conditions were significantly correlated (i.e., for distribution of the neural pattern similarity across participants, see **Fig. S4**; r (mean ± SD): = 0.151 ± 0.417, Fisher’ z (mean ± SD) = 0.223 ± 0.570, t(36) = 2.38, p = 0.023, p(permutation) = 0.012).

#### 3.3.2 Functional connectivity varies across different dilemmas in vmPFC-related network (GLM-PPI)

By using the conjunction findings of vmPFC as the seed region (see **SI: fMRI study methods** for details), we observed that the functional connectivity between vmPFC and clusters including the dmPFC extending to the supplementary motor area (whole-brain level corrected), and the left TPJ (peak MNI coordinates: -56/-62/22, t(36) = 3.53, p(SVC-FWE) = 0.026) was significantly higher in the *self-bad* cause dilemma when making immoral decisions (vs. *charity-bad cause*; see **Fig. 5**; also see **Table S2** for details of PPI results in each dilemma separately).

**Fig. 5.**
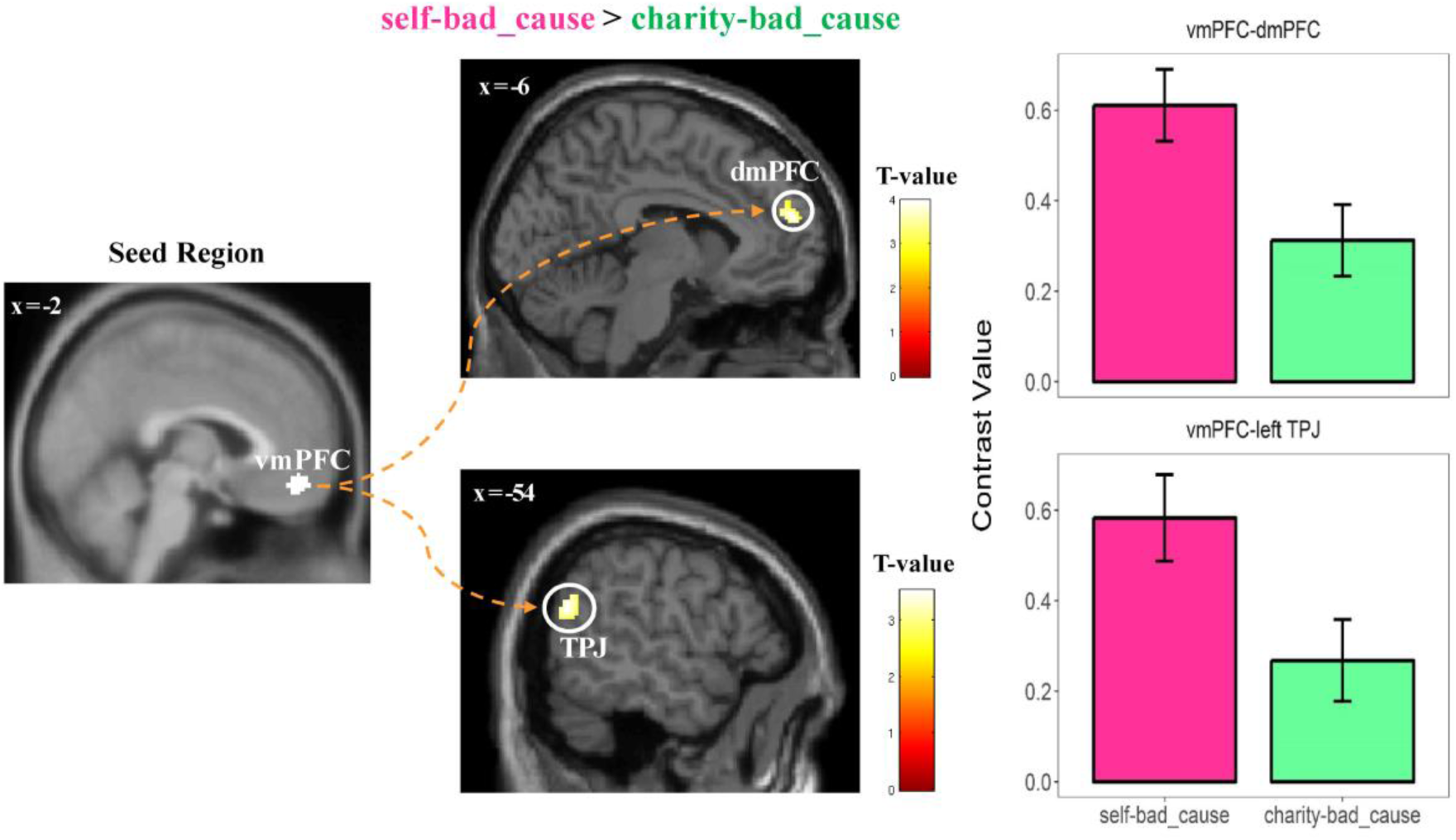
Reduced connectivity between vmPFC and mentalizing network (i.e., dorsomedial prefrontal cortex, dmPFC; left temporoparietal junction, TPJ) during decision-making period in the *charity-bad_cause* dilemma (vs. *self-bad_cause*). To visualize these connectivity results, we extracted the contrast value of the PPI regressor in both regions (i.e., masks are defined by the intersection between the activated cluster and the independent ROI; dmPFC: center MNI: x/y/z = -1/56/24; left TPJ: center MNI: x/y/z = -53/-59/20; spheres with a radius of 9mm) in each dilemma. No further statistics were performed to avoid of double dipping. Display threshold: p < 0.001 uncorrected at the voxel-level with k = 100.

#### 3.3.3 Neural correlates of single attributes (i.e., monetary gains and moral cost; GLM2)

For the completeness of the analyses, we also examined the neural correlates of single attributes in each dilemma separately. In the *self-bad cause* dilemma, we found a positive modulation of personal monetary gains on the neural signals in bilateral ventral striatum and medial prefrontal areas from mid-cingulate cortex to the ventral part of the anterior cingulate cortex (ACC), whereas a negative modulation of moral costs was observed in activity of bilateral inferior parietal lobules (IPL) and left orbitofrontal cortex. In the *charity-bad cause* dilemma, we observed a positive modulation of money with increasing donations to the charity on the activities in the dorsal part of the ACC, the SMA and the left IPL, whereas a negative modulation of moral costs was observed in the dorsomedial prefrontal cortex (dmPFC) and the right IPL (see **Fig. S5** and **Table S3** for details of all activated regions). No difference on the parametric effect of moral cost was observed between the two dilemmas. The conjunction analysis on the positive modulation of the monetary gains in two dilemmas and that on the negative modulation of the moral costs did not reveal any significant brain region.

#### 3.3.4 Individual differences in moral preference modulate on context-dependent choice-specific decision-relevant neural activation (GLM3)

The right dlPFC (i.e., peak MNI coordinates: 40/36/18, t(31) = 4.52, p(SVC-FWE) = 0.003) was the only brain region showing a positive correlation between moral preferences and the dilemma × decision interaction (i.e., contrast comparing *accept* vs. *reject* in the interaction between *self-bad* cause vs. *charity-bad* cause dilemma). To better understand this correlation, we extracted contrast values in right dlPFC (defined by an independent mask; see **Methods** for details) and ran correlation analyses between the right dlPFC signals during acceptance (vs. reject) choice and moral preference in each dilemma separately, using the lmrob function in the “rubustbase” package in R (Rousseeuw *et al*., 2015), which rules out the effect of outlier data points. As a result, the moral preference positively correlated with the right dlPFC activity during acceptance (vs. reject) in the *self-bad cause* dilemma (robust correlation: r (95% CI) = 0.31 (0.14, 0.49), t(31) = 3.64, p < 0.001) whereas a trend-to-significant negative correlation was observed in the *charity-bad cause* dilemma (r (95% CI) = −0.29 (−0.61, 0.02), t(31) = −1.93, p = 0.063; see **Fig. 6**).

**Fig. 6.**
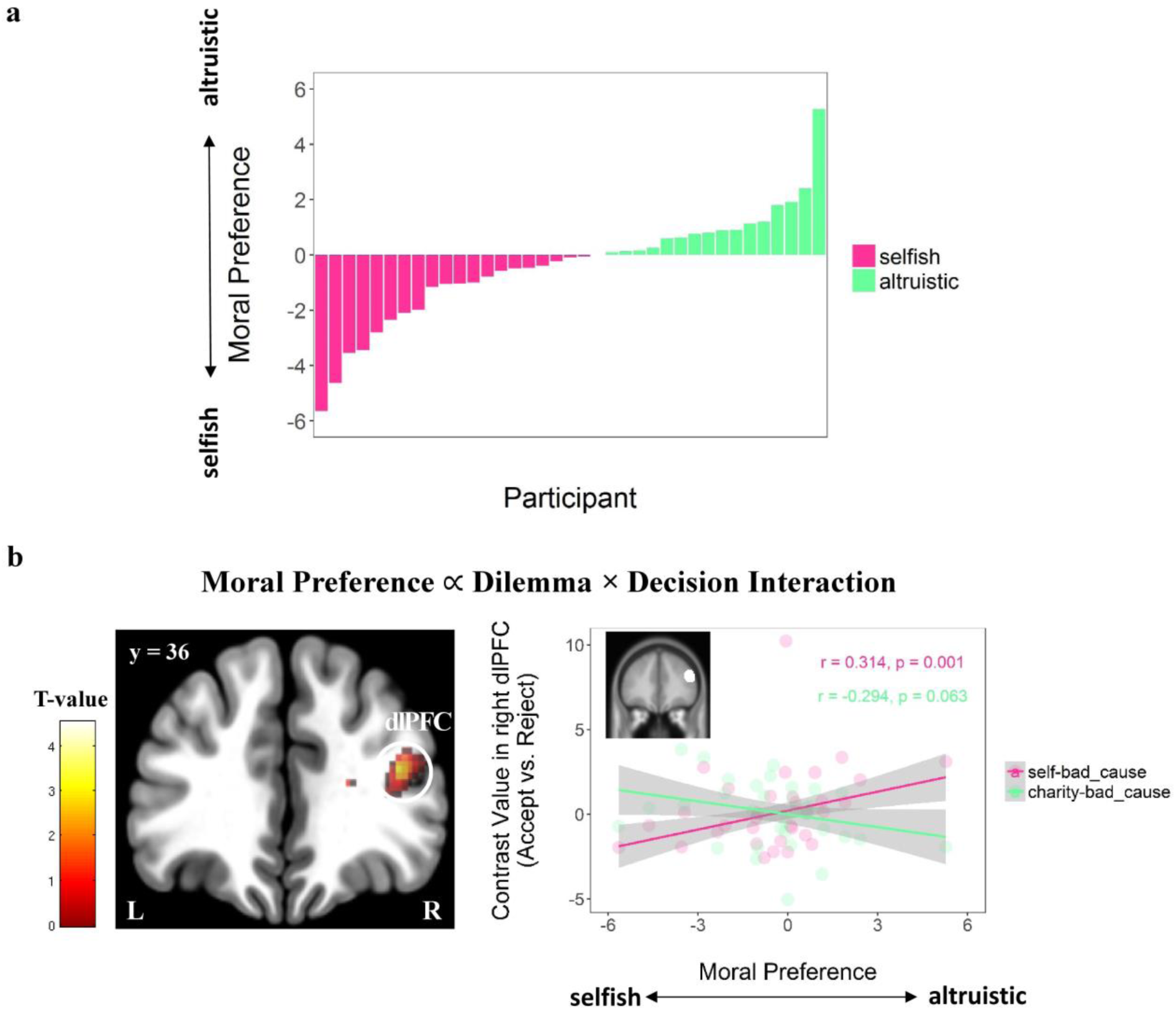
Individual differences in moral preference modulate on context-dependent choice-specific decision-relevant neural activation. (a) Histogram of the distribution of moral preference across participants. The higher (i.e., more positive) the moral preference is, the more the participant weight on the relative gain for the charity compared with themselves (i.e., more altruistic). (b) Individual difference in moral preference modulates the decision × dilemma interactive effect on the right dorsolateral prefrontal cortex (dlPFC; GLM3; N = 33). To unpack the modulation effect, we extracted the contrast value averaged within the right dlPFC (masked by an independent ROI; center MNI: x/y/z = 46/36/24; spheres with a radius of 9mm) from accept vs. reject contrasts in each dilemma, showing that more altruistic participants tend to elicit higher activity in the right dlPFC when they accept (vs. reject) the offer in the *self-bad_cause* dilemma but lower activity when making the same choice in the *charity-bad_cause* dilemma. Robust correlation was used to rule out the effect of outlier. Lines represent the robust linear fits. Shaded areas represent the 95% confidence interval. Display threshold: p < 0.001 uncorrected at the voxel-level with k = 50.

## 4. Discussion

It is well established that people modify their immoral behaviors depending on exactly who will benefit (Lewis *et al*., 2012; Garrett *et al*., 2016), but the mechanisms allowing this plasticity in immorality have yet to be described. To address this issue, we designed a novel task in which participants in the scanner were asked to make a series of decisions benefiting either themselves or a charity while simultaneously yielding an immoral consequence in *both* conditions. Even though participants reported that they felt it was more morally inappropriate to accept an immoral offer to benefit themselves rather than a charity, they nevertheless accepted self-serving immoral offers more frequently and more quickly than those benefitted a preferred charity. These findings are in accordance with previous studies revealing that people exhibit self-serving bias in moral judgment (Bocian and Wojciszke, 2014) and decision-making (Engel, 2011). Moreover, these results agree with the proposal of moral license, such that displaying altruism via another outlet (i.e., accepting the immoral offers to benefit a charity) would license people to behave more selfishly by accepting more self-serving immoral offers (Merritt *et al*., 2010). Corroborating this interpretation, we observed a strong inter-individual positive correlation between the acceptance rates of the two types of moral dilemma (involving self or charity as beneficiaries). This compensatory mechanism was also reflected in the fact that participants accepting a self-serving immoral offer in a previous trial were more likely to accept subsequent other-serving immoral offers. These results suggest that individuals may justify their immoral behaviors (i.e., earning morally-tainted profits for themselves) by performing alternative altruistic acts (i.e., accepting an immoral offer to benefit a charity), which presumably contributes to a positive self-concept (i.e., the way people view and perceive themselves; Aronson, 1969; Baumeister, 1998).

By adopting a computational approach, we further offer a mechanistic explanation for the behavioral differences underlying these immoral decisions. The winning computational model among those tested assumed the subjective value of an immoral offer as a linear summation of the weighed monetary gain and moral cost. Such weights varied in different dilemmas: although participants weighed the value of monetary gains for themselves or the charity along similar lines, they differed with respect to how they treated moral cost. That is, participants weighed the moral cost more negatively in dilemma benefiting a charity (*vs. self)*, thus explaining why participants accepted offers less frequently in this condition. Notably, our winning computational model also extends a previously reported model accounting for the prosocial choices engaging a tradeoff between personal gains and donations to a charity (Lopez-Persem et al., 2017) to the immoral domain. The present findings also provide a computational account and direct support for the theory of self-concept maintenance (Aronson, 1969; Mazar *et al*., 2008), which, until this study, has only been tested without computational modeling. This theory proposes that individuals generally value morality (e.g., honesty), and therefore want to maintain this aspect of their self-concept (Greenwald, 1980; Sanitioso *et al*., 1990). Here, if a person fails to comply with his internal standards for morality by profiting himself with ill-gotten gains, he will need to negatively update his self-concept by decreasing the weight on the moral cost (Mazar *et al*., 2008).

At the brain system level, we showed that valuation of immoral offers was associated with relative subjective value computation in the vmPFC, integrating monetary benefits and moral cost regardless of the beneficiaries (self or charity). This finding is consistent with a unified neural representation of value in which the vmPFC is regarded as the key hub (Bartra *et al*., 2013; Clithero and Rangel, 2013). Furthermore, our RSA analysis directly demonstrated that the neural patterns of vmPFC during value computation identified in the *self-bad cause* dilemma is similar to the one in the *charity-bad cause* dilemma. The vmPFC is recruited in the integration of positive (e.g., monetary reward) and negative stimuli (e.g., electrical shock) during value-based decision making processes in non-social contexts (Basten *et al*., 2010; Park *et al*., 2011). Previous studies have indicated a similar role for the vmPFC in moral valuation and decision-making (Moll *et al*., 2005; Greene, 2014). However, these studies did not investigate whether the same valuation regions were engaged for immoral decisions regardless of the beneficiaries. We observed that the functional coupling between the vmPFC and mentalizing regions (i.e., dmPFC together with the TPJ) (Schaafsma *et al*., 2015), was enhanced when making immoral decisions benefiting oneself (as compared to the charity). This differential functional coupling according to the beneficiary of the decision parallels our behavioral findings of a higher acceptance rate for immoral decisions benefiting oneself as compared to a charity. The brain valuation system is known to work together with the mentalizing network during social decision-making (e.g., strategic interactive decisions) (Hampton *et al*., 2008; Hill *et al*., 2017). In particular, the TPJ is engaged when weighing self-interest with other regarding preferences (Strombach *et al*., 2015) and it has been shown to be causally necessary to signal the moral conflict between personal profits and moral costs (Obeso *et al*., 2018). In the present study, the enhanced functional coupling between the vmPFC and mentalizing nodes (i.e., dmPFC and TPJ) in the *self-bad cause* vs. *charity-bad cause* dilemma, reflects that the cross talk between the valuation and the mentalizing network is more sensitive to the signal involving a conflict between personal interests (as compared to the welfare of others) and moral cost. This finding is also interesting in light of recent reports that the mentalizing network supports a key role for social interactions involving oneself as compared to situations in which the participant is an observer rather than an interactive participant (Redcay and Schilbach, 2019).

Furthermore, we investigated the link between choice-specific brain activity and individual differences in moral preference (i.e., an index measuring the extent of behavioral changes in different dilemmas) and how this relationship is influenced by the beneficiary. We found large inter-individual differences in moral preferences at the behavioral level, in line with previous studies (Crockett *et al*., 2014; Crockett *et al*., 2017; Yin *et al*., 2017). Investigation of the relationship between this behavioral effect and brain activation revealed, as predicted, that the dlPFC activity was associated with inter-individual differences in moral preference and that this relationship also depended upon the beneficiary of the immoral choice. *Post-hoc* analyses showed that this interaction was a consequence of moral preference increasing with enhanced dlPFC activity in the *self-bad cause* dilemma when participants accepted (vs. rejected) the offer, whereas the opposite relationship occurred in the *charity-bad cause* dilemma. The dlPFC has been shown to impact on a variety of social and moral decisions, such as fairness (Knoch *et al*., 2006; Spitzer *et al*., 2007; Ruff *et al*., 2013), generosity (Hutcherson *et al*., 2015) and dishonesty (Greene and Paxton, 2009; Maréchal *et al*., 2017), and has also been linked to individual differences in moral behavior (Crockett *et al*., 2017). In a recent theoretical framework, the lPFC, including but not limited to the dorsal part, was proposed to act as a flexible guide in the pursuit of moral goals, which depend on the interaction between context and a specific individual (Carlson and Crockett, 2018). In the present study, participants with increasing positive scores in moral preference can be considered as more altruistic while those with decreasing negative scores can be considered to be more selfish. Critically, a negative correlation between this task-based score and the Machiavellianism trait further confirmed the underlying rationale and external validity of our measure of moral preference. Thus, as reflected by stronger dlPFC signals associated with acceptance of *self-bad cause* dilemmas (vs. *charity-bad cause*), altruistic people (i.e., those with higher positive moral preference scores) have to overcome a stronger subjective moral cost when accepting offers that profit themselves. Therefore, our findings provide empirical evidence in support of the hypothesis that dlPFC engagement is key to explaining inter-subject variability in solving conflicts between self-interest and moral cost. In line with the present finding, a recent fMRI investigation found that participants are willing to trade their moral values (i.e., similar to the moral cost in the present study), in exchange for personal profit, and this effect is accompanied by decision value computation engaging the lateral PFC (Qu *et al*., 2019).

Despite that obtaining personal gains *via* an immoral approach is universally forbidden by moral rules, people still break such moral rules by trading-off self-interest and moral values to maintain a positive self-image. However, the process of moral decision-making can be highly flexible across different contexts, meaning that the balance point between different cost-benefit tradeoffs is not fixed, but context-dependent. Our findings identify the neurocomputational mechanisms and the brain circuitry underlying this flexible immoral behavior to provide a mechanistic understanding of the neurobiological architecture encompassing value computation and integration of contextual information. In particular, an overall value signal is constructed from independent attributes and this integration between single attributes is computed in the vmPFC (computing the decision value) regardless of the beneficiary. In turn, stronger functional connectivity between the vmPFC and components of the mentalizing network (i.e., dmPFC and TPJ) during the dilemma weigh up concerns for the bad cause and oneself as compared to the bad cause and the charity. This represents the neural signature of flexible moral behavior depending upon the beneficiary of the immoral action. Our results also shed light on the question of whether moral decisions rely on the same valuation circuitry engaged during value-based decision making or requires a specialized set of brain regions. From a broader perspective, these findings may have societal implications because the way we value immorality fundamentally affects our behaviors and further shapes the functioning of our societies.

## Acknowledgments

This research has benefited from the financial support of IDEXLYON from Université de Lyon (project INDEPTH) within the Programme Investissements d’Avenir (ANR-16-IDEX-0005) and of the LABEX CORTEX (ANR-11-LABX-0042) of Université de Lyon, within the program Investissements d’Avenir (ANR-11-IDEX-007) operated by the French National Research Agency. J-C.D. was funded by the grant “ANR-NSF CRCNS ‘SOCIAL_POMDP’ n°16-NEUC”. Q.C. was funded by the Program for National Science Foundation of China (31470995). We would like to thank Si-ying Li, Yan-er Su, and Fang-jie Xu for their assistance during data collection, Dr. Chen Hu, Dr. Lei Zhang, Dr. Xiao-xue Gao as well as Dr. Martin Hebart for their help on data analyses, Dr. Chun-liang Feng for comments on an earlier draft.

## Supporting Information

### SI: Association selection

#### Procedure

An independent group of participants (N = 30) attended the rating task for selecting the associations used for the fMRI study. 16 different associations were used for the rating task. Besides those associations used in the fMRI study (i.e., 4 charities and 4 associations supporting the rights to own guns or hunting), there were another 4 charities and 4 associations supporting the legalization of soft drugs (see **vignettes and logos** below; also see **Fig. S6a**). Participants read vignettes (with the logo) which described the goal of each association. Vignettes were presented in a pseudo-random fashion.

Below each vignette, participants were asked to complete a series of questions by indicating degrees on a Likert rating scale, including 1) degree of familiarity (0 = “not at all”, 10 = “very much”), 2) degree of liking (−10 = “not at all” or “very negative”, 0 = “no preference”, 10 = “very much” or “very positive”), 3) degree of moral acceptance (−10 = “not at all”, 0 = “no preference”, 10 = “very much”), 4) degree of approval for its goal (−10 = “not at all”, 0 = “no preference”, 10 = “very much”), 5) amount of (hypothetical) donation (from 0 to 10; in CNY) and 6) degree of moral conflict while donating (0 = “not at all”, 10 = “very much”).

A rating summary of all associations in all questions by all participants is listed in **Table S4**. To induce stronger morally negative feelings and control the familiarity difference between charities and bad causes, we finally decided to use associations in support of gun ownership or hunting as the morally bad causes and charities with lower familiarity for the current fMRI study (see **Table S4** and **Fig. S6b**).

To justify our selection, we compared the rating scores in all questions between the charities and morally bad causes used in the current fMRI study. Neither the charities nor the morally bad causes were familiar to participants, indicated by the low average scores of familiarity (i.e., less than 2 on a 0-10 Likert scale; mean ± SD: charity: 1.3 ± 1.4; morally bad cause: 0.5 ± 0.8). As expected, participants donated much more (charity vs. bad cause [mean ± SD; same below]: 6.9 ± 2.2 vs. 0.6 ± 1.2; b (95% confidence interval (CI)) = 6.34 (5.76, 6.92), SE = 0.28, t(6) = 22.37, p < 0.001, Cohen’d = 18.3) and showed significantly higher levels of liking (7.1 ± 1.7 vs. -5.0 ± 2.9; b (95% CI) = 12.13 (10.83, 13.42), SE = 0.67, t(6) = 18.12, p < 0.001, Cohen’d = 14.8), moral acceptance (8.2 ± 1.6 vs. -3.7 ± 3.1; b (95% CI) = 11.84 (9.70, 13.98), SE = 1.11, t(6) = 10.70, p < 0.001, Cohen’d = 8.73), approval (8.0 ± 1.7 vs. -4.7 ± 3.1; b (95% CI) = 12.71 (11.21, 14.20), SE = 0.77, t(6) = 16.54, p < 0.001, Cohen’d = 13.50) and less moral conflict with the charities than bad causes (0.9 ± 1.7 vs. 6.6 ± 2.5; b (95% CI) = −5.70 (−6.41, −4.99), SE = 0.35, t(6) = −16.21, p < 0.001, Cohen’d = −13.23).

#### Vignettes and Logos

##### Charity

###### Charities familiar to local participants (CF)

**International Committee of the Red Cross (ICRC**; official website: https://www.icrc.org) was founded 1863 and is a humanitarian institution based in Geneva, Switzerland. It is one of the most widely recognized organizations in the world, having won three Nobel Peace Prizes in 1917, 1944, and 1963. ICRC aims to protect victims of international and internal armed conflicts. Such victims include war wounded, prisoners, refugees, civilians, and other non-combatants.

(Information source: https://en.wikipedia.org/wiki/International_Committee_of_the_Red_Cross)

**The United Nations Children’s Fund (UNICEF**; official website: https://www.unicef.org/) was founded in 1946 and is a United Nations (UN) program headquartered in New York City that provides humanitarian and developmental assistance to children and mothers in developing countries. UNICEF programs emphasize developing community-level services to promote the health and well-being of children. UNICEF was awarded the Nobel Peace Prize in 1965 and the Prince of Asturias Award of Concord in 2006.

(Information source: https://en.wikipedia.org/wiki/UNICEF)

**SOS Children’s Villages** (official website: http://www.sos-childrensvillages.org/) was founded in 1949 and is an independent, non-governmental international development organization which has been working to meet the needs and protect the interests and rights of children. The organization’s work focuses on abandoned, destitute and orphaned children requiring family-based child care. Children are supported to recover from being emotionally traumatized and to avoid the real danger of being isolated, abused, exploited and deprived of their rights.

(Information source: https://en.wikipedia.org/wiki/SOS_Children%27s_Villages)

**World Wide Fund for Nature (WWF**; official website: https://en.wikipedia.org/wiki/World_Wide_Fund_for_Nature) was founded in 1961 and is an international non-governmental organization, working in the field of the wilderness preservation, and the reduction of humanity’s footprint on the environment. It is the world’s largest conservation organization with over five million supporters worldwide. The group’s mission is “to stop the degradation of the planet’s natural environment and to build a future in which humans live in harmony with nature.” Currently, much of its work concentrates on the conservation of oceans and coasts, forests, and freshwater ecosystems.

(Information source: https://en.wikipedia.org/wiki/World_Wide_Fund_for_Nature)

###### Charities unfamiliar to local participants (CUF; finally used in the pilot and the fMRI study)

**Oxford Committee for Famine Relief** (OXFAM; official website: https://www.oxfam.org/) was founded in 1942 in Oxford, and is an international confederation of charitable organizations focused on the alleviation of global poverty. OXFAM’s programs address the structural causes of poverty and related injustice and work primarily through local accountable organizations, seeking to enhance their effectiveness. OXFAM’s stated goal is to help people directly when local capacity is insufficient or inappropriate for OXFAM’s purposes and to assist in the development of structures which directly benefit people facing the realities of poverty and injustice. (Information source: https://en.wikipedia.org/wiki/Oxfam)

**Save the Children** (also known as the Save the Children Fund; official website: https://www.savethechildren.net/) was founded in 1919 in London, and is an international non-governmental organization that promotes children’s rights, provides relief and helps support children in developing countries. Save the Children uses a holistic approach to help us achieve more for children, and to use resources in an efficient and sustainable way.

(Information source: https://en.wikipedia.org/wiki/Save_the_Children; https://www.savethechildren.net/about-us)

**First Aid Africa** (FAA; official website: http://www.firstaidafrica.org/) was founded in 2008 in Edinburg and is a humanitarian charity that works in rural parts of southeastern Africa to provide sustainable equipment and education in first aid. FAA explains that a small amount of medical knowledge and equipment’ can make a difference. Volunteers and students receive some training before traveling to Africa to teach first aid and survival skills in settings such as local communities, schools, orphanages, and villages.

(Information source: https://en.wikipedia.org/wiki/First_Aid_Africa)

**Oceana** (official website: http://oceana.org/) was found in 2001, Washington and is the largest international ocean conservation and advocacy organization. Oceana works to protect and restore the world’s oceans through targeted policy campaigns. Oceana bases its policy campaign goals on science to achieve concrete and measurable results through targeted campaigns that combine policy, advocacy, science, law, media, and public pressure to prevent the collapse of fish populations, marine mammals and other sea life caused by industrial fishing and pollution. (Information source: https://en.wikipedia.org/wiki/Oceana_(non-profit_group))

##### Morally Bad Cause

###### Drug Legalization Associations (D)

**Law Enforcement Action Partnership** (**LEAP**; formerly Law Enforcement Against Prohibition; official website: https://lawenforcementactionpartnership.org/) was founded in 2002 and is a non-profit, international, educational organization comprising former and current police officers, government agents and other law enforcement agents who oppose the current War on Drugs. In January 2017, while reaffirming their commitment to ending the War on Drugs, LEAP became the Law Enforcement Action Partnership in order to advocate for solutions across a broader range of drug policy and criminal justice issues.

(Information source: https://en.wikipedia.org/wiki/Law_Enforcement_Action_Partnership; https://www.dinafem.org/en/blog/cannabis-marijuana-legalization-groups/)

**European Coalition for Just and Effective Drug Policies (ENCOD**; official website: http://www.encod.org/info/-English-en-.html) was founded in 1993 and is a European non-governmental organization which brings European citizens together who believe that drug prohibition is an immoral and insane policy. Since 1994 they have been working to advocate more just and effective drugs control policies, which include an integrated solution for all problems related to the global drugs phenomenon.

(Information source: https://en.wikipedia.org/wiki/European_Coalition_for_Just_and_Effective_Drug_Policies; https://www.dinafem.org/en/blog/cannabis-marijuana-legalization-groups/)

**Marijuana Policy Project (MPP**; official website: https://www.mpp.org/) was founded in 1995 and is the largest organization working solely on marijuana policy reform in the United States. Its stated aims include: (1) increase public support for non-punitive, non-coercive marijuana policies and (2) change state laws to reduce or eliminate penalties for the medical and non-medical use of marijuana. MPP believes the greatest harm associated with marijuana is a prison, so their focus is on removing criminal penalties for marijuana use.

(Information source: https://en.wikipedia.org/wiki/Marijuana_Policy_Project; https://www.dinafem.org/en/blog/cannabis-marijuana-legalization-groups/)

**National Organization for the Reform of Marijuana Laws (NORML**; official website: http://norml.org/) was founded in 1970 and is an American non-profit organization whose aim is to move public opinion sufficiently to achieve the legalization of non-medical marijuana in the United States so that the responsible use of cannabis by adults is no longer subject to penalty. NORML supports the removal of all criminal penalties for the private possession and responsible use of marijuana by adults, including the cultivation for personal use, and the casual nonprofit transfers of small amounts.

(Information source: https://en.wikipedia.org/wiki/National_Organization_for_the_Reform_of_Marijuana_Laws)

##### Morally Bad Cause

###### Gun/Hunting Rights Advocacy Associations (GH; finally used in the pilot and the fMRI study)

**National Rifle Association of America (NRA**; official website: https://home.nra.org/) is an American nonprofit organization which advocates for gun rights. Founded in 1871 to advance rifle marksmanship, the modern NRA continues to teach firearm competency and safety. NRA has been criticized by newspaper editorial boards, gun control and gun rights advocacy groups, political commentators, and politicians. For instance, a Washington Post/ABC News poll in January 2013 showed that only 36 percent of Americans had a favorable opinion of NRA leadership.

(Information source: https://en.wikipedia.org/wiki/National_Rifle_Association)

**The European Federation of Associations for Hunting and Conservation of the EU (FACE**; organization which is pro-hunting and pro-gun. FACE protested a 2016 proposal by the European Union to revise the EU’s firearm regulations, saying it would ban muzzle-loading weapons. In 2016 FACE opposed an EU proposal to ban the import of hunting trophies from certain African countries.

(Information source: https://en.wikipedia.org/wiki/Federation_of_Associations_for_Hunting_and_Conservation_of_the_EU)

**The Society for Liberal Weapons Rights (ProTell**; official website: https://www.protell.ch/de/) is a Swiss gun-rights advocacy group based in Bern, Switzerland. The association was founded in 1978 with the purpose of defending the right of law-abiding citizens to carry arms, and is opposed to any restrictions in this regard. ProTell was one of the principal opponents to the federal popular initiative “For the protection against gun violence”, brought to a referendum on February 13, 2011. The initiative was broadly rejected by the voters.

(Information source: https://en.wikipedia.org/wiki/ProTell)

**Safari Club International (SCI**; official website: https://www.safariclub.org/) is an international organization composed of hunters dedicated to protecting the freedom to hunt and promoting wildlife conservation. SCI has been criticized for supporting the hunting of endangered African antelope species at fenced “game” ranches in Texas and Florida and for giving awards for hunting leopards, elephants, lions, rhinos and buffalo in Africa.

(Information source: https://en.wikipedia.org/wiki/Safari_Club_International)

### SI: Pilot behavioral study

#### Methods

Thirty undergraduates or graduate students (15 females; mean age: 20.9 ± 1.7 years, ranging from 18 to 25 years) were recruited via online fliers for the pilot behavioral study.

The procedure and behavioral paradigm was same as the fMRI study, except the following: 1) besides the original payoff matrix (i.e., monetary gain: moral cost = 1:4), we adopted a balanced matrix with the payoffs ranging from 1 to 8 (in increments of 1; unit: CNY) for both parties involved in the dilemma (i.e., monetary gain: moral cost = 1:1), thereby producing 64 different payoff combinations as offers. As a consequence, the pilot study was extended to 256 trials in total (i.e., 64 trials for each context of moral dilemma displaying offers produced by different payoff matrices); 2) To maximize the differential effect induced by two types of payoff matrix, we adopted a mixed design so that offers from the same payoff matrix were presented in a block (i.e., one block for original and balanced matrix respectively) with the different dilemmas randomly intermixed in-between. The order of block was counterbalanced across participants; 3) To reduce the total duration of this experiment, we used a 500ms inter-trial interval (ITI) showing a cross fixation.

We analyzed the data with a similar approach as the fMRI study except that we added the predictors (i.e., main effect or interaction terms) relevant to matrix type (i.e., dummy variable; reference-level: original matrix) in the regression analyses.

#### Results

All these associations were selected by participants at least once (see **Fig. S7a**). Participants were familiar to none of the selected associations, indicated by the low average scores of familiarity (i.e., less than 3 on a 0-10 Likert scale; mean ± SD: charity: 2.80 ± 3.10; bad cause: 1.63 ± 2.27). Moreover, they rated the pre-selected charity positively (mean ± SD (95% CI): 7.83 ± 2.00 (7.09, 8.58); t(29) = 21.44, p < 0.001, Cohen’s d = 3.91) whereas they regarded the pre-selected bad cause negatively (mean ± SD (95% CI); −7.87 ± 2.54 (−8.82, −6.92); t(29) = −16.95, p < 0.001, Cohen’s d = 3.09; see **Fig. S7b**).

We found a significant matrix × context interaction effect on the choice data (Odds Ratio = 4.33, b (95% CI) = 1.46 (1.19, 1.74), SE = 0.14, χ^2^(1) = 106.63, p < 0.001) after controlling the monetary gain and the moral cost. Splitting the data in terms of the matrix type, we found that participants were less likely to accept the offer in the *charity-bad_cause* dilemma (vs. *self-bad_cause* dilemma: accept rate: 26.4 ± 25.3 % vs. 40.6 ± 37.4 %; Odds Ratio = 0.24, b (95% CI) = −1.44 (−1.66, −1.22), SE = 0.11, χ^2^(1) = 170.77, p < 0.001) only when the original payoff matrix was adopted. No difference in acceptance rate between two contexts of moral dilemma was observed when the balanced payoff matrix was used (62.1 ± 21.8 % vs. 62.3 ± 28.5 %; Odds Ratio = 0.98, b (95% CI) = −0.02 (−0.25, 0.21), SE = 0.12, χ^2^(1) = 0.03, p = 0.861; see **Fig. S7c**; also see **Table S5** for details of model output).

For the decision time (DT), we first did the log-transformation due to its non-normal distribution (Anderson-Darling normality test: A = 219.3, p < 0.001). Regressions on log-transformed DT (logDT) revealed a three-way interaction (b (95% CI) = 0.10 (0.03, 0.16), SE = 0.03, t(7643) = 2.93, p = 0.003, Cohen’s d = 0.07; see **Table S6** for descriptive summary of DT and logDT). To unpack the interaction effect, we ran similar regression analyses with matrix, context and their interaction as fixed-effect predictors for trials with acceptance and rejection decisions separately. For acceptance trials, we observed a significant two-way interaction on logDT (b (95% CI) = −0.12 (−0.16, −0.07), SE = 0.02, t(3644) = −4.80, p < 0.001, Cohen’d = −0.16). Again, we did the simple effect analyses by splitting the accept trials into two parts depending on matrix type, finding that such two-way interaction was mainly driven by the stronger effect of dilemma type in prolonging logDT in the *charity-bad_cause* dilemma (vs. *self-bad_cause* dilemma) when offers were chosen from the original matrix (b (95% CI) = 0.15 (0.11, 0.19), SE = 0.02, t(1265) = 7.41, p < 0.001, Cohen’d = 0.42) than the balanced matrix (b (95% CI) = 0.04 (0.01, 0.06), SE = 0.01, t(2359) = 2.74, p = 0.006, Cohen’d = 0.21). For reject trials, no two-way interaction effect was detected (b (95% CI) = 0.02 (−0.02, 0.06), SE = 0.02, t = 0.970, p = 0.332, Cohen’d = 0.03). We only found that people rejected more slowly in the block displaying offers from balanced (vs. original) payoff regardless of dilemmas (b (95% CI) = 0.07 (0.04, 0.11), SE = 0.02, t(3970) = 3.97, p < 0.001, Cohen’d = 0.13; see **Fig. S8**; also see **Table S7** for details of model output).

#### Behavioral Analyses

All behavioral analyses were conducted using R (http://www.r-project.org/) and relevant packages (R Core Team, 2014). All reported p values are two-tailed and p < 0.05 was considered statistically significant. Data visualization were performed via “ggplot2” package (Wickham H 2016).

Regarding the choice data, we performed a repeated mixed-effect logistic regression on the decision of choosing the “accept” option by the glmer function in “lme4” package (Bates D et al. 2013), with dilemma (dummy variable; reference level: *self-bad_cause* dilemma; same below) and payoffs for both parties involved in each dilemma (i.e., the monetary gain and the moral cost; mean-centered continuous variable; same below) as the fixed-effect predictors. In addition, we included the following random-effect factors allowing varying intercept across participants. For the statistical inference on each predictor, we performed the Type II Wald chi-square test on the model fits by using the Anova function in “car” package (Fox J et al. 2016), and reported the odds ratio as relevant effect size.

For decision time (DT), we first did a log-transformation due to its non-normal distribution (Anderson-Darling normality test: A = 91.90, p < 0.001) and then performed a mixed-effect linear regression on the log-transformed DT by the lmer function in “lme4” package, with decision (dummy variable; reference level: accept), dilemma, decision × dilemma, as well as payoffs for both beneficiaries as the fixed-effect predictors. Random-effect factors were specified in the same way as above. Similar analyses were also performed on the post-scanning rating except that dilemma was added as the only fixed-effect predictor. We followed the procedure recommended by Luke (2017) to obtain the statistics for each predictor by applying the Satterthwaite approximations on the restricted maximum likelihood model (REML) fit via the “lmerTest” package (Luke SG 2017). In addition, we computed the Cohen’d of each predictor via the “EMAtools” package (Kleiman E 2017), which provided the effect size measure specially for the mixed-effect regressions. For likeness ratings of the selected associations, we compared whether the ratings significantly differed from 0 in each type of selected associations (i.e., charity or morally bad causes) respectively by the one-sample T-test, and computed the Cohen’s d as effect size.

### Supporting Figures

**Fig. S1.**
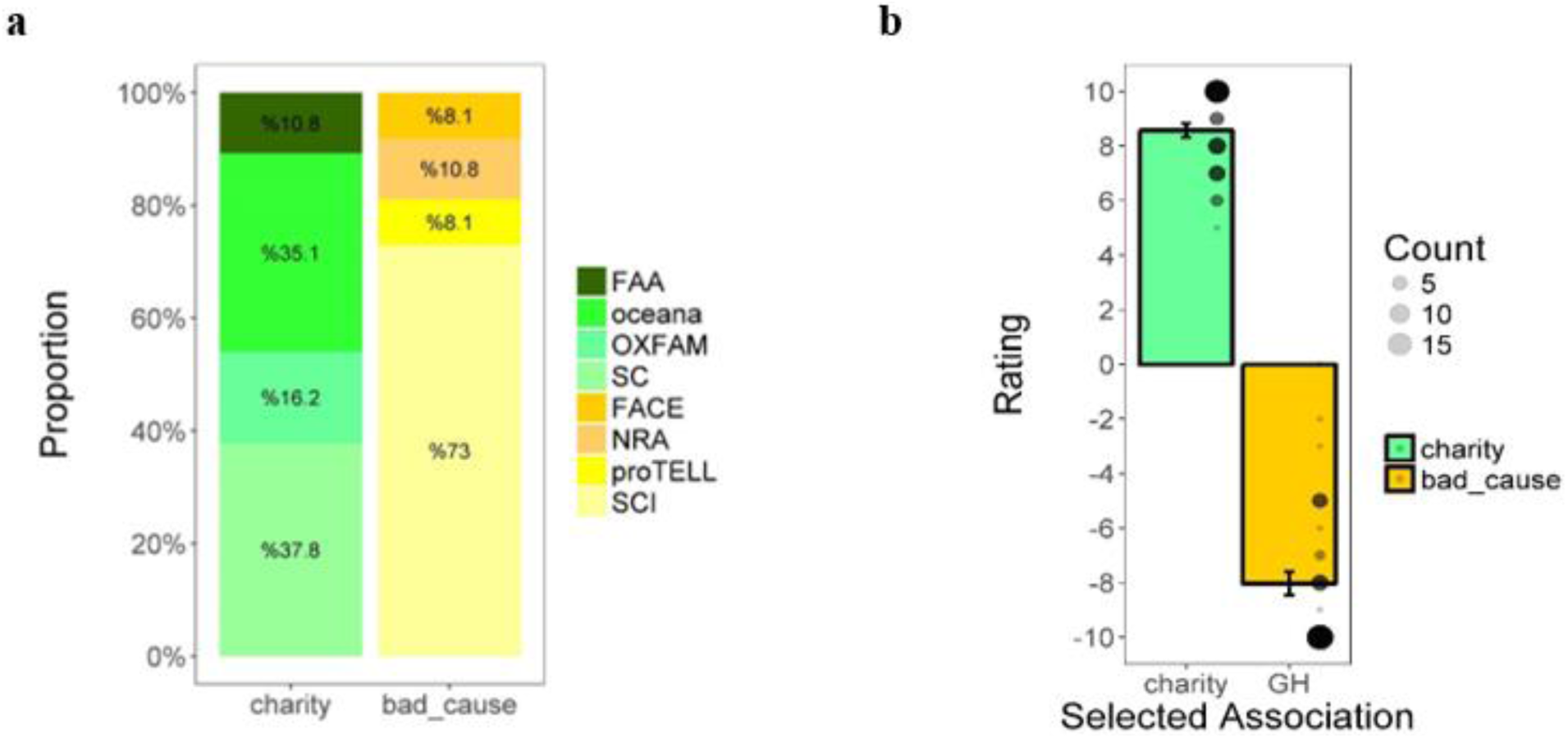
(a) Summary for the charity and morally bad cause selected by all participants and (b) mean rating on liking (−10 – 10; −10 = dislike very much, 10 = like very much) for the selected charities and morally bad causes in the current fMRI study. The black dot refers to the individual ratings; the size of the dot indicates the numbers of participants with the same rating.

**Fig. S2.**
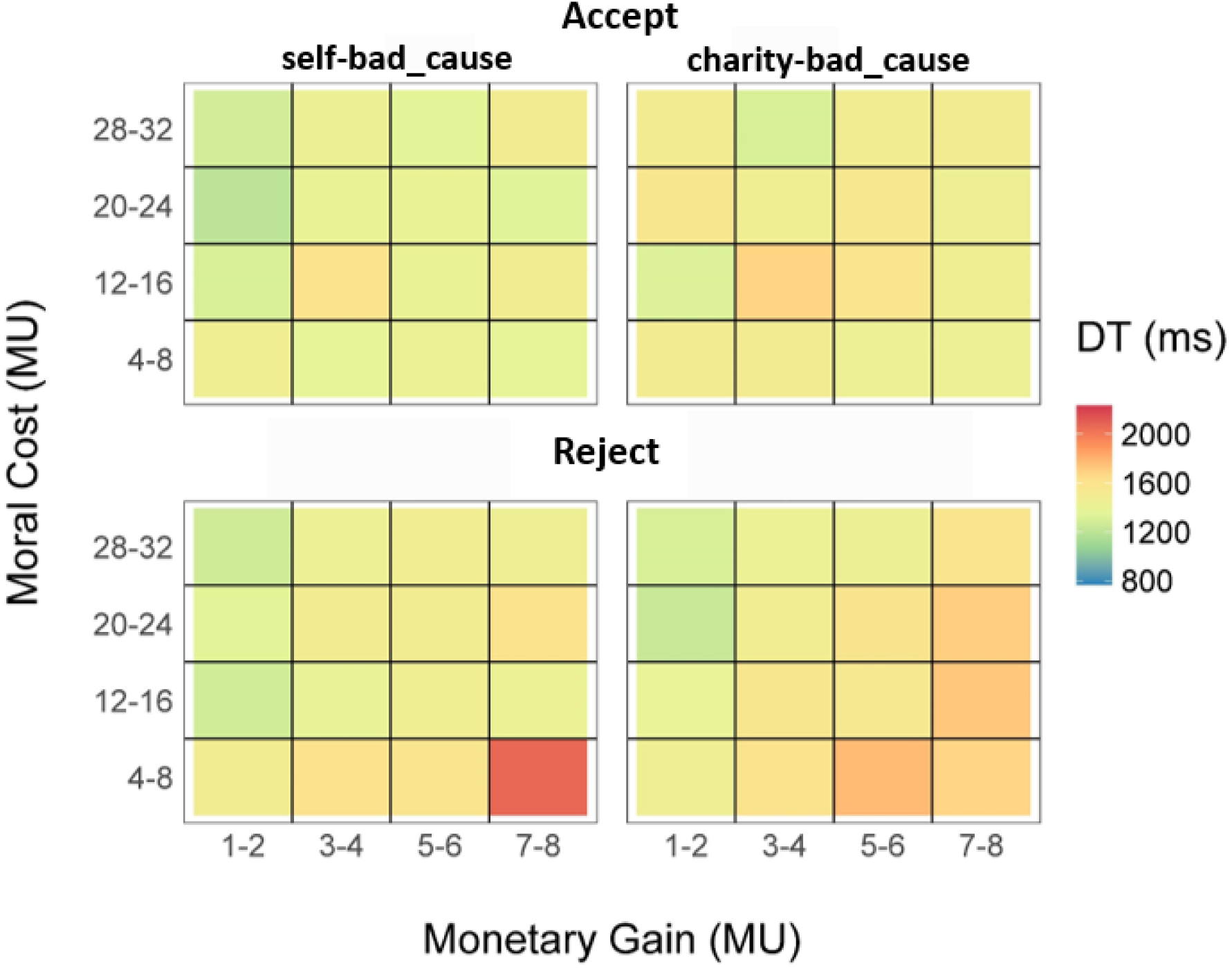
Heat map of the mean decision time (DT) as a function of the monetary gain and the moral cost in two dilemmas of the current fMRI study. Data were collapsed into 4-by-4 matrices only for a better visualization.

**Fig. S3.**
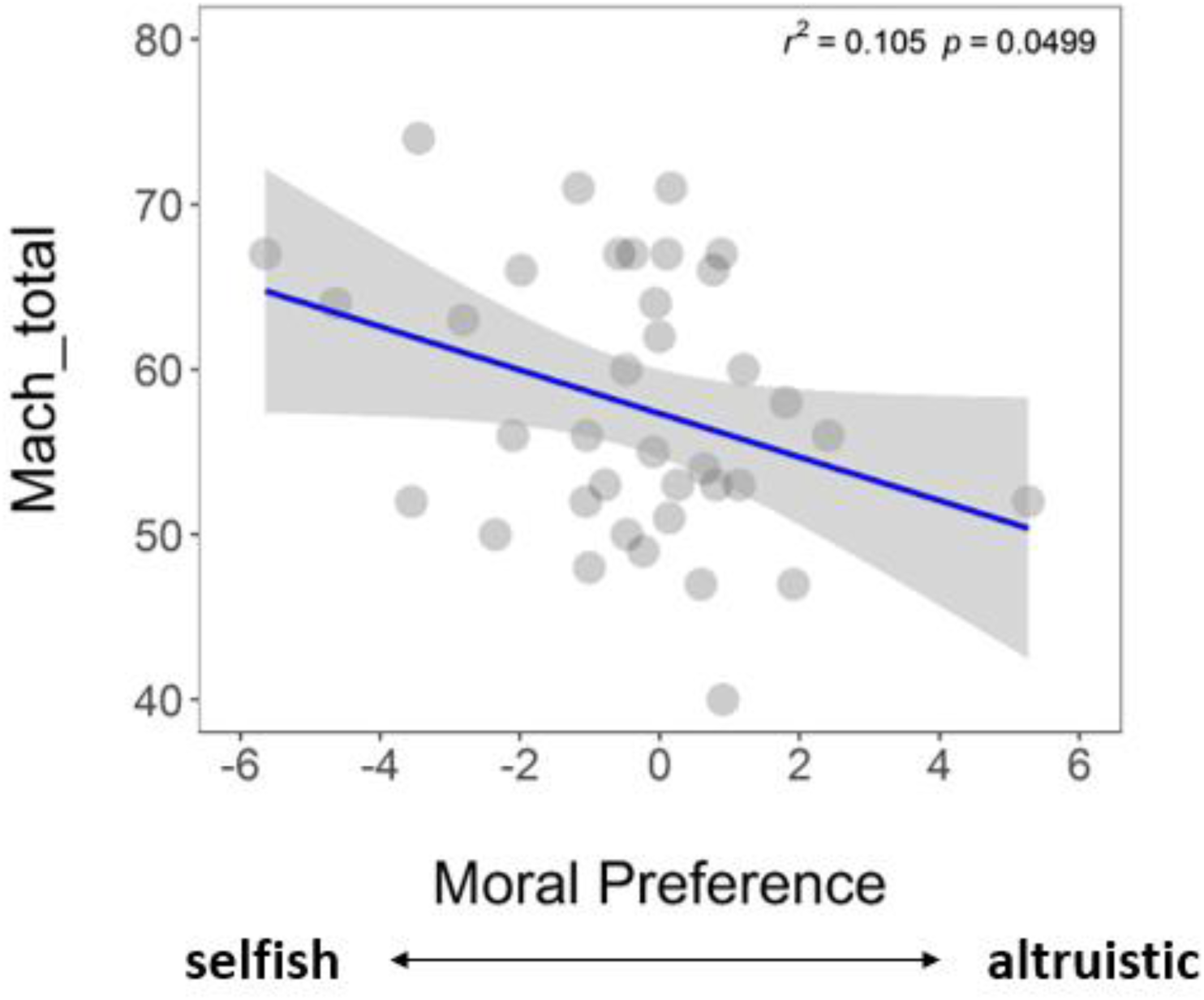
Negative correlation between the moral preference and the Machiavellian score. The higher the Machiavellian score is, the higher degree that a person agrees with the idea of pursuing personal gain via immoral approaches. Each dot represents the data of a single participant.

**Fig. S4.**
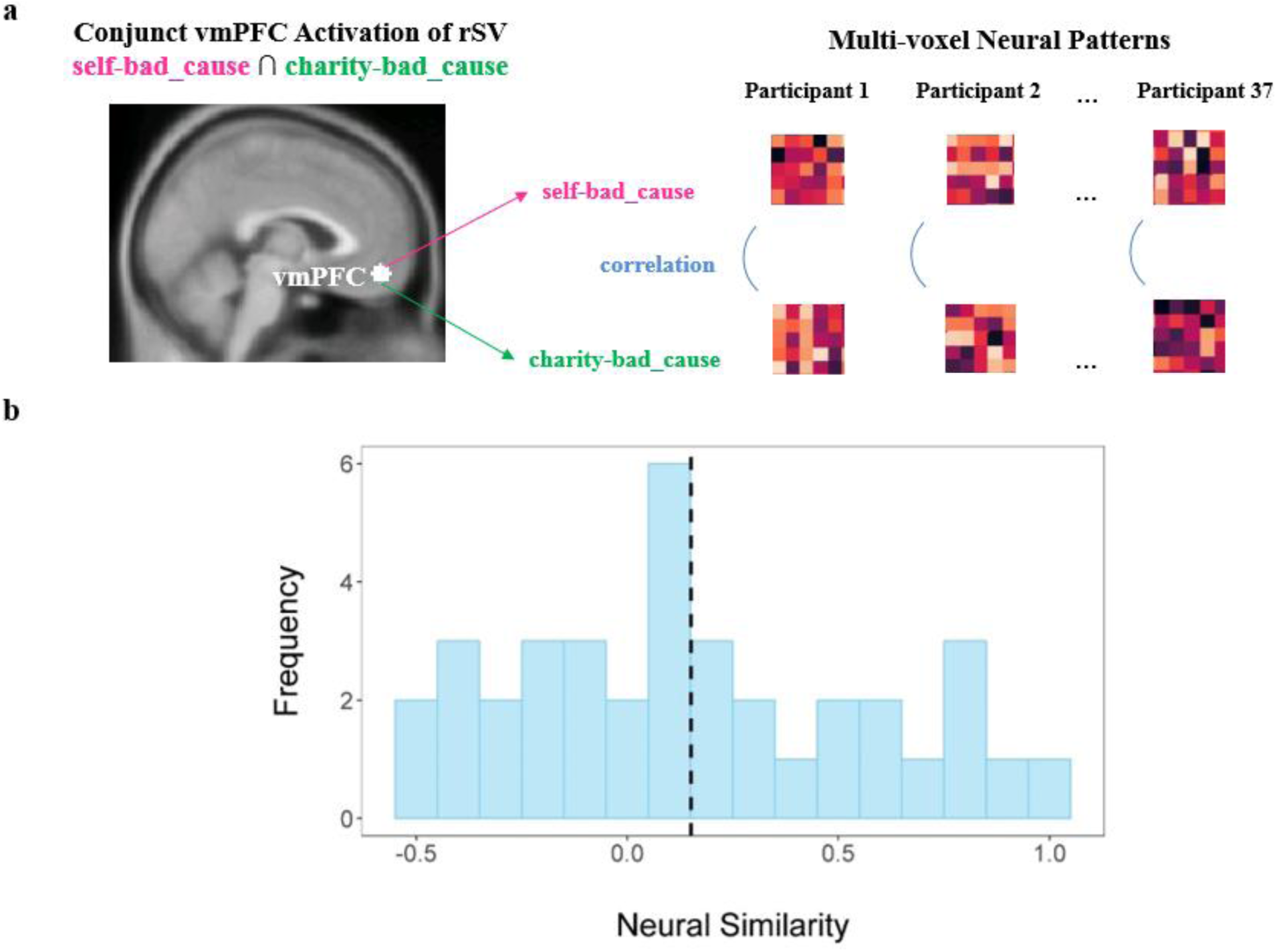
Procedure and results of representational similarity analysis (RSA). (a) For each participant, we first defined the vmPFC mask based on the conjunction activation in GLM1 (peak MNI: -2/48/-14; a sphere with a radius of 6mm). Then we extracted the multi-voxel neural patterns (i.e., those heat maps; only for illustration) within the vmPFC mask from the contrast image characterizing the parametric effect of relative subjective value (SV) in each dilemma. Next we computed the dissimilarity between these neural patterns in two dilemmas and obtained the correlation coefficients (i.e., similarity) using one minus dissimilarity. For statistical analysis, all correlation coefficients were transformed to Fisher’s z value. (b) Histogram of the distribution of neural similarity between across all participants. The vertical dashed line refers to the mean of the neural similarity.

**Fig. S5.**
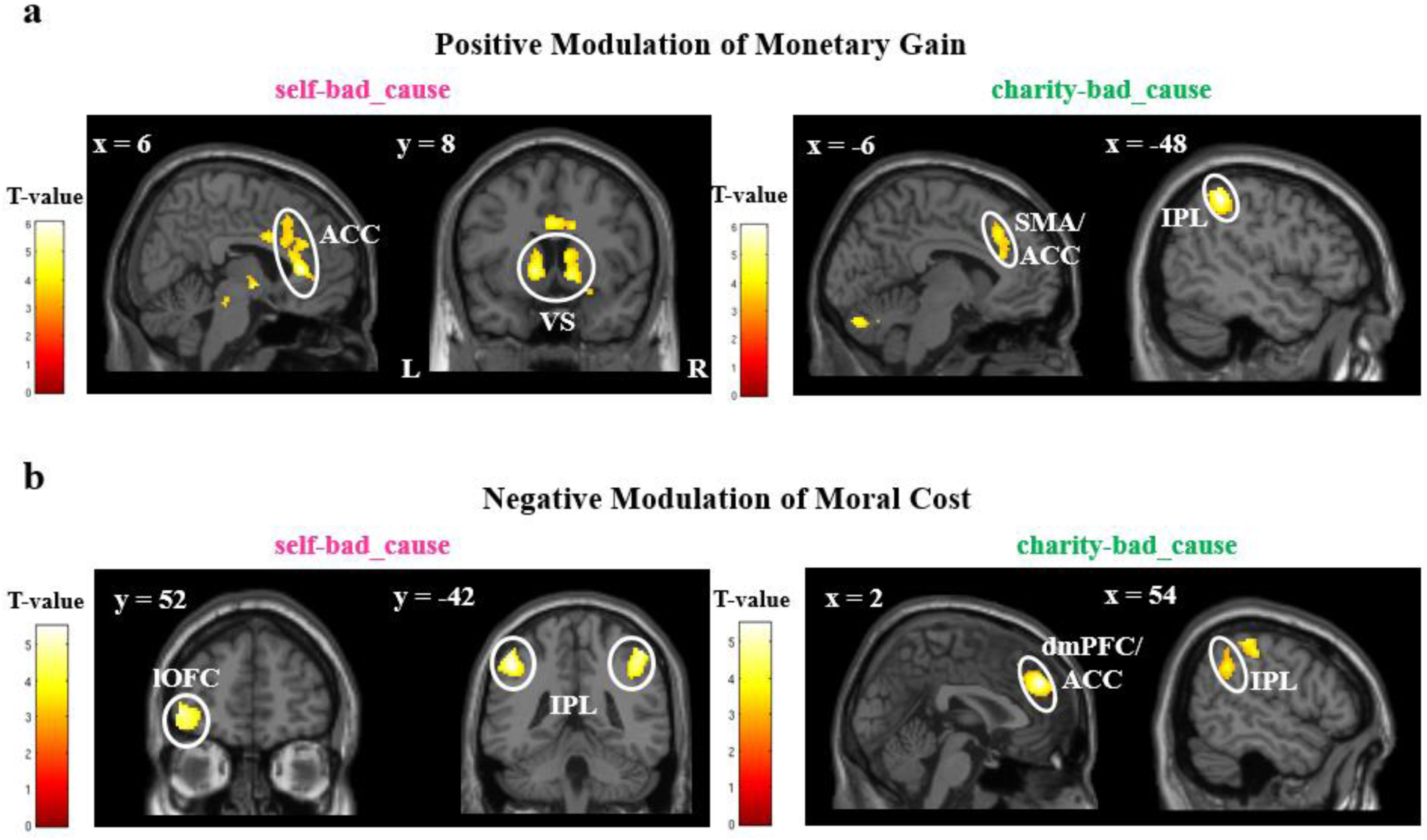
Neural correlates of single attributes. (a) Positive modulation of the monetary gain (i.e., payoff for oneself and the charity) in each dilemma. (b) Negative modulation of the moral cost (i.e., benefits for the bad cause) in each dilemma. Abbreviation: ACC = anterior cingulate cortex; dmPFC = dorsomedial prefrontal cortex; SMA = supplementary motor area; IPL = inferior parietal lobule; lOFC = lateral orbitofrontal cortex; VS = ventral striatum; Display threshold: p < 0.001 uncorrected at the voxel-level with k = 200.

**Fig. S6.**
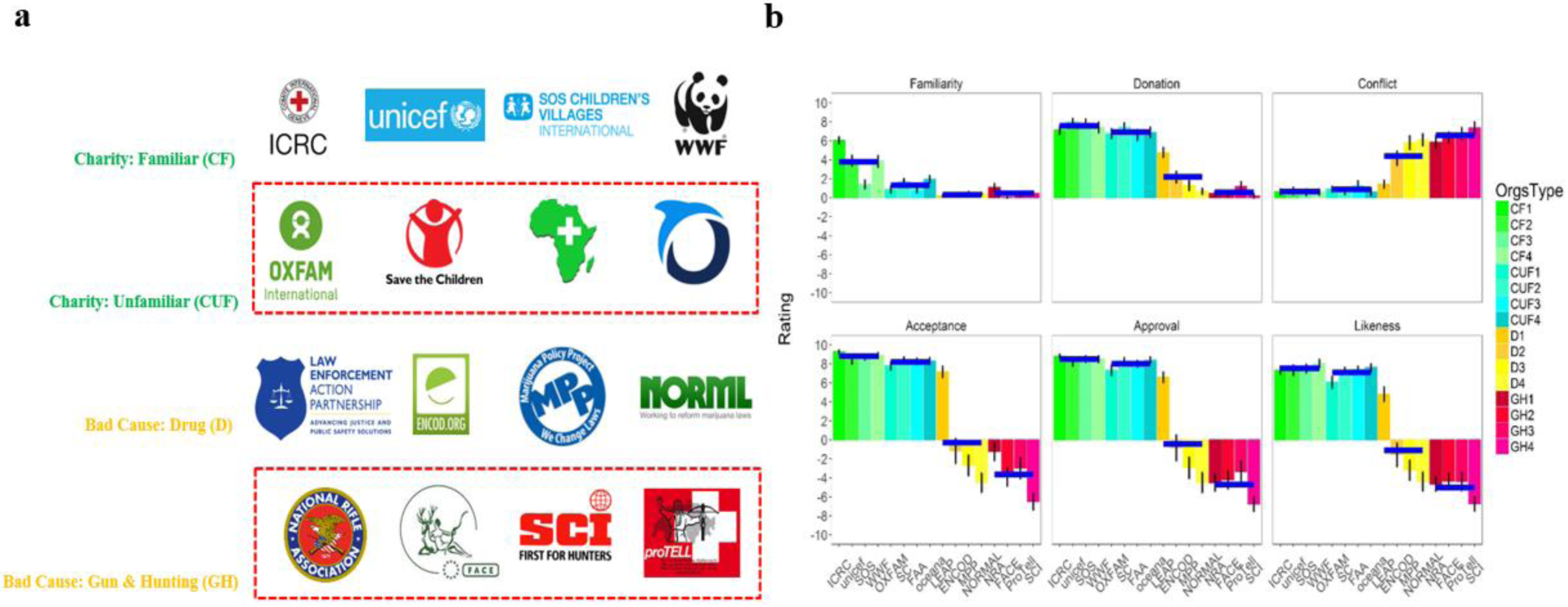
(a) All logos of associations used for selection. Those charities and morally bad causes are marked by the red frame we finally used for the pilot. (b) Mean ratings on individual associations as well as categories according to six dimensions in the questionnaire.

**Fig. S7.**
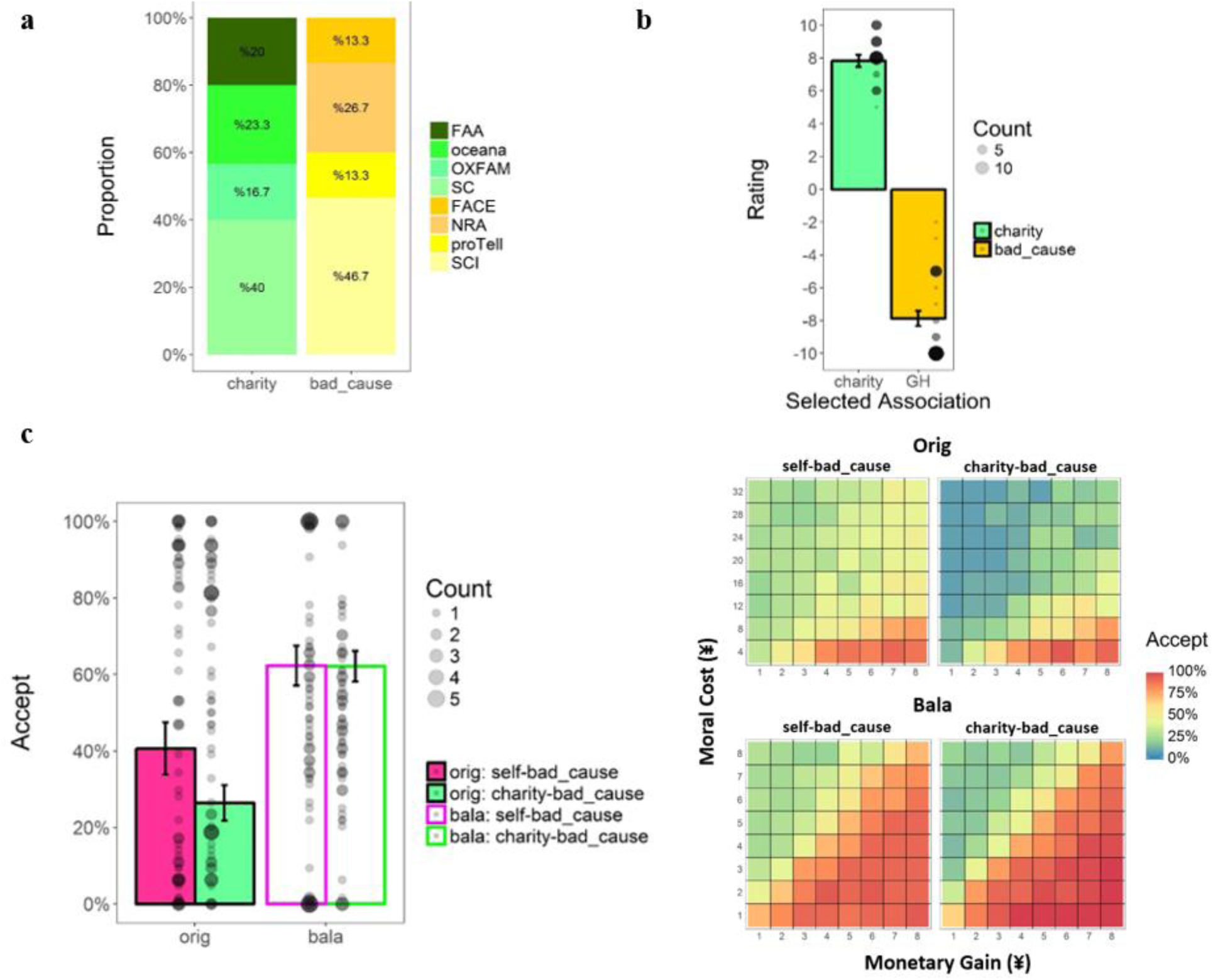
Results of the pilot behavioral study. (a) Summary for the charity and the morally bad cause selected by all participants. (b) Mean rating on liking (−10 – 10; −10 = dislike very much, 10 = like very much) for the selected charity and morally bad cause. The black dot refers to the individual ratings; the size of the dot indicates the numbers of participants with the same rating. (c) Left panel: the mean acceptance rate in each dilemma with each payoff matrix; Right panel: the heat map of the mean acceptance rate (%) at each payoff amount for each beneficiary involved in both dilemmas. Abbreviations: orig= original payoff matrix (i.e., monetary gain: moral cost = 1:4); bala = balanced payoff matrix (i.e., monetary gain: moral cost = 1:1).

**Fig. S8.**
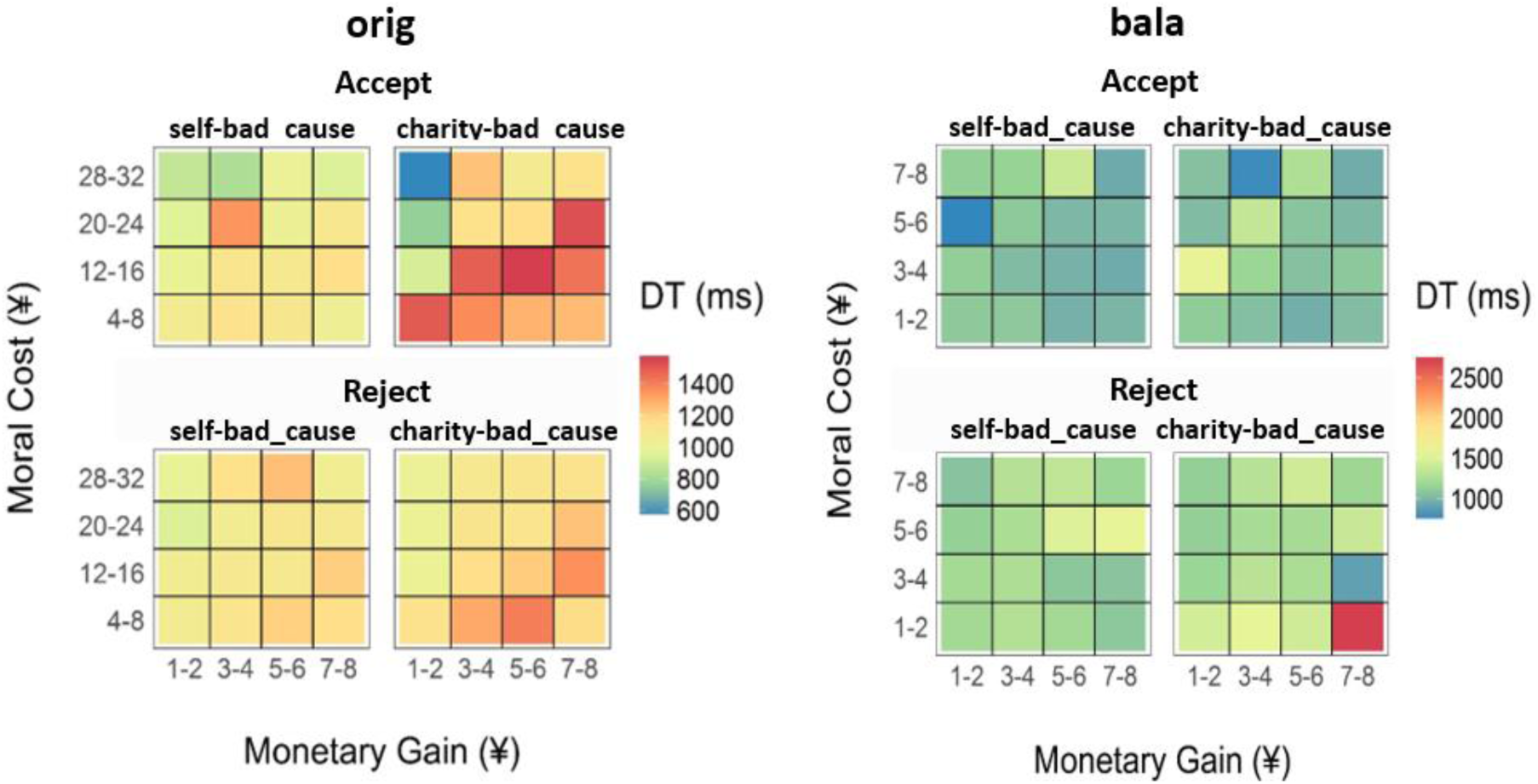
Heat map of the mean decision time (DT) for specific choice as a function of the monetary gain and the moral cost in each dilemma, separated by different payoff matrix, in the pilot behavioral study. Data were collapsed into 4-by-4 matrices only for a better visualization. Abbreviations: orig= original payoff matrix (i.e., monetary gain: moral cost = 1:4); bala = balanced payoff matrix (i.e., monetary gain: moral cost = 1:1).

### Supporting Tables

**Table S1.** Regions encoding the relative subjective value (rSV) during decision-making process in each dilemma (N = 37, GLM1)

| Brain Region | Hemisph<br>ere | Cluster<br>Size | MNI |  |  | BA | T-value | p(cl-FWE) |
| --- | --- | --- | --- | --- | --- | --- | --- | --- |
|  |  |  | x | y | z |  |  |  |
| ***self-bad_cause*** |  |  |  |  |  |  |  |  |
| **rSV: positive modulation** |  |  |  |  |  |  |  |  |
| PCG/MFG | L | 258 | -28 | -8 | 46 | 6 | 4.49 | 0.046 |
| PCG/MFG | R | 282 | 30 | -4 | 42 | 6 | 4.67 | 0.034 |
| FuG/LG/PHG/IOG | L | 671 | -36 | -56 | -12 | 19/37 | 5.42 | 0.001 |
| FuG/LG/PHG/<br>MOG/IOG/MTG | R | 1652 | 26 | -70 | 2 | 18/19/20/<br>36/37 | 5.47 | < 0.001 |

Note: Regions shown here met the Family-Wise Error corrected cluster-level (cl-FWE) threshold of $p < 0.05$ , with an uncorrected voxel-level threshold of $p < 0.001$ as the cluster-defining threshold. No significant region was found in positive or negative modulation of rSV in the *charity-bad\_cause* dilemma under this threshold. Coordinates shown here were based on Montreal Neurological Institute (MNI) coordinate system. Abbreviations: L: left, R: right, B: bilateral, BA: Brodmann Area; MFG: middle frontal gyrus, PCG: pre-central gyrus, MTG: middle temporal gyrus, MOG: middle occipital gyrus, IOG: inferior occipital gyrus, FG: fusiform gyrus, LG: lingual gyrus, PHG: parahippocampal gyrus.

**Table S2.** Regions showing enhanced functional connectivity with vmPFC in different dilemmas (i.e., *PPI: self-bad_cause* vs. *implicit baseline*; *PPI: charity-bad_cause* vs. *implicit baseline*; N = 37, gPPI-GLM)

| Regions | Hemisphere | Cluster Size | MNI |  |  | BA | T-value | p(cl-FWE) |
| --- | --- | --- | --- | --- | --- | --- | --- | --- |
|  |  |  | x | y | z |  |  |  |
| **PPI: *self-bad_cause* vs. *implicit baseline*** |  |  |  |  |  |  |  |  |
| vmPFC/MeFG/dmPFC/ACC | B | 2022 | -2 | 38 | -12 | 9/10/11/24/32 | 11.76 | < 0.001 |
| SFG | R | 53 | 22 | 34 | 46 | 8 | 6.20 | < 0.001 |
| PCC/Prec | B | 757 | -4 | -62 | 30 | 7/23/30/31 | 8.30 | < 0.001 |
| MTG/ITG | L | 60 | -56 | -12 | -20 | 21 | 7.06 | < 0.001 |
| ITG | R | 150 | 60 | -8 | -20 | 21 | 7.77 | < 0.001 |
| IFG/STG | L | 67 | -32 | 16 | -22 | 38/47 | 7.69 | < 0.001 |
| AG/SmG/STG/MTG | L | 304 | -46 | -66 | 28 | 39 | 7.41 | < 0.001 |
| AG/SmG/STG/MTG | R | 331 | 60 | -52 | 28 | 39/40 | 7.86 | < 0.001 |
| Cerebellum | L | 57 | -10 | -52 | -36 |  | 6.53 | < 0.001 |
| **PPI: *charity-bad_cause* vs. *implicit baseline*** |  |  |  |  |  |  |  |  |
| vmPFC/ACC | B | 766 | -4 | 50 | -12 | 10/11/32 | 11.40 | < 0.001 |
| PCC/Prec |  | 310 | -8 | -58 | 10 | 23/30 | 6.82 | < 0.001 |

Note: A FWE-corrected voxel-level threshold of $p < 0.05$ with $k = 50$ was adopted. Coordinates shown here were based on Montreal Neurological Institute (MNI) coordinate system. Abbreviations: p(cl-FWE): cluster-level Family-Wise Error corrected threshold; L: left, R: right, B: bilateral, BA: Brodmann Area; ACC: anterior cingulate cortex, AG: angular gyrus, dmPFC: dorsomedial prefrontal cortex, MeFG: medial frontal gyrus, MTG, middle temporal gyrus, SFG: superior frontal gyrus, ITG: inferior temporal gyrus, SmG: supramarginal gyrus, PCC: posterior cingulate cortex, STG: superior temporal gyrus, vmPFC: ventromedial prefrontal cortex, Prec: precuneus.

**Table S3.** Regions encoding single attributes (i.e., monetary gain and moral cost) during decision-making process in each dilemma (N = 37, GLM2)

| Brain Region | Hemisphere | Cluster Size | MNI |  |  | BA | T-value | p(cl-FWE) |
| --- | --- | --- | --- | --- | --- | --- | --- | --- |
|  |  |  | x | y | z |  |  |  |
| ***self-bad_cause*** |  |  |  |  |  |  |  |  |
| **monetary gain:** |  |  |  |  |  |  |  |  |
| **positive modulation** |  |  |  |  |  |  |  |  |
| VS (Caud/Put/Pal)/Tha | L | 471 | -12 | 0 | -4 |  | 6.06 | 0.005 |
| VS (Caud/Put/Pal)/IFG | R | 574 | 14 | 4 | -4 | 47 | 5.03 | 0.002 |
| ACC/MCC/SMA/MeFG/SFG | B | 1277 | 6 | 32 | 8 | 8/24/32 | 5.55 | < 0.001 |
| **moral cost:** |  |  |  |  |  |  |  |  |
| **negative modulation** |  |  |  |  |  |  |  |  |
| IFG/MFG | L | 413 | -54 | 40 | -4 | 10/11/47 | 5.09 | 0.004 |
| MFG/IFG | R | 451 | 36 | 26 | 40 | 8/9 | 4.95 | 0.003 |
| IPL/AG/SmG/PoCG | L | 607 | -48 | -42 | 46 | 40 | 5.51 | < 0.001 |
| IPL/SPL/AG/SmG/STG/PoCG | R | 1572 | 48 | -46 | 56 | 7/39/40 | 5.04 | < 0.001 |
| ***charity-bad_cause*** |  |  |  |  |  |  |  |  |
| **monetary gain:** |  |  |  |  |  |  |  |  |
| **positive modulation** |  |  |  |  |  |  |  |  |
| MeFG/SMA/ACC/MCC | B | 445 | -10 | 30 | 38 | 6/8/9/32 | 4.45 | 0.001 |
| IPL | R | 300 | -48 | -42 | 50 | 40 | 6.11 | 0.010 |
| Cerebellum | L | 349 | -32 | -68 | -30 |  | 5.05 | 0.005 |
| Cerebellum | R | 675 | 36 | -58 | -32 |  | 5.00 | < 0.001 |
| **moral cost:** |  |  |  |  |  |  |  |  |
| **negative modulation** |  |  |  |  |  |  |  |  |
| MeFG/SFG/ACC/MCC | B | 1037 | 2 | 46 | 34 | 6/8/9/32 | 6.45 | < 0.001 |
| SFG/MeFG/MFG | R | 330 | 12 | 38 | 52 | 8 | 4.67 | 0.013 |
| IPL/SmG/PoCG/AG | R | 439 | 58 | -26 | 44 | 40 | 5.46 | 0.003 |

Note: Regions shown here met the Family-Wise Error corrected cluster-level (cl-FWE) threshold of $p < 0.05$ , with an uncorrected voxel-level threshold of $p < 0.001$ as the cluster-defining threshold. No significant region was found in the negative modulation of monetary gain and the positive modulation of moral cost, in each dilemma respectively. Coordinates shown here were based on Montreal Neurological Institute (MNI) coordinate system. Abbreviations: L: left, R: right, B: bilateral, BA: Brodmann Area; ACC: anterior cingulate cortex, MCC: mid-cingulate cortex, MFG: middle frontal gyrus, MeFG: medial frontal gyrus, IFG: inferior frontal gyrus, SFG: superior frontal gyrus, SMA: supplementary motor area, AG: angular gyrus, PoCG: postcentral gyrus, SmG: supramarginal gyrus, IPL: inferior parietal lobule, STG: superior temporal gyrus, VS: ventral striatum; Caud: caudate, Put: putamen, Pal: pallidum.

**Table S4.** Summary of average rating (mean ± SD) in the rating task for association selection (N = 30)

|  |  | Familiarity | Likeness | Acceptance | Approval | Donation | Conflict |
| --- | --- | --- | --- | --- | --- | --- | --- |
| CF | ICRC | 6.0 ± 2.2 | 7.3 ± 2.3 | 9.3 ± 1.0 | 8.8 ± 1.2 | 7.1 ± 2.6 | 0.6 ± 1.4 |
| | unicef | 3.9 ± 3.0 | 7.3 ± 2.8 | 8.4 ± 2.4 | 8.2 ± 2.6 | 8.0 ± 2.1 | 0.7 ± 1.9 |
| | SOS | 1.4 ± 2.4 | 7.5 ± 2.4 | 8.7 ± 1.5 | 8.5 ± 1.8 | 7.9 ± 2.4 | 0.6 ± 1.9 |
| | WWF | 3.8 ± 3.3 | 8.0 ± 2.5 | 8.9 ± 1.5 | 8.5 ± 2.0 | 7.4 ± 3.1 | 0.6 ± 1.8 |
| | pooled | 3.8 ± 2.0 | 7.6 ± 1.8 | 8.8 ± 1.3 | 8.5 ± 1.5 | 7.6 ± 2.2 | 0.6 ± 1.6 |
| CUF | **OXFAM** | **0.9 ± 1.9** | **6.1 ± 3.3** | **7.8 ± 2.4** | **7.3 ± 3.0** | **6.7 ± 2.8** | **0.9 ± 1.6** |
|  | **SC** | **1.7 ± 2.0** | **7.3 ± 2.0** | **8.4 ± 2.0** | **8.3 ± 2.0** | **7.4 ± 2.4** | **0.7 ± 2.0** |
|  | **FAA** | **0.8 ± 1.7** | **7.4 ± 1.6** | **8.3 ± 1.7** | **8.0 ± 1.9** | **6.6 ± 3.0** | **1.3 ± 2.8** |
|  | **Oceana** | **2.0 ± 2.1** | **7.6 ± 2.2** | **8.3 ± 1.9** | **8.4 ± 1.8** | **6.9 ± 2.9** | **0.6 ± 2.0** |
|  | **pooled** | **1.3 ± 1.4** | **7.1 ± 1.7** | **8.2 ± 1.6** | **8.0 ± 1.7** | **6.9 ± 2.2** | **0.9 ± 1.7** |
| D | LEAP | 0.2 ± 0.6 | 4.8 ± 4.2 | 7.2 ± 3.1 | 6.6 ± 3.0 | 4.8 ± 2.7 | 1.4 ± 2.3 |
| | ENCOD | 0.3 ± 0.8 | -1.6 ± 6.7 | -1.2 ± 6.9 | -0.8 ± 7.2 | 2.2 ± 3.3 | 4.2 ± 4.0 |
| | MPP | 0.4 ± 1.5 | -3.2 ± 5.7 | -2.7 ± 5.9 | -2.9 ± 6.1 | 1.3 ± 2.9 | 5.8 ± 3.8 |
| | NORML | 0.4 ± 0.9 | -4.4 ± 5.4 | -4.5 ± 5.6 | -4.5 ± 5.7 | 0.7 ± 1.7 | 6.1 ± 3.5 |
| | pooled | 0.3 ± 0.7 | -1.1 ± 3.7 | -0.3 ± 3.6 | -0.4 ± 3.8 | 2.2 ± 2.0 | 4.4 ± 2.3 |
| GH | **NRA** | **1.1 ± 2.1** | **-4.7 ± 4.0** | **-1.2 ± 5.0** | **-4.5 ± 4.7** | **0.5 ± 1.5** | **5.9 ± 3.5** |
|  | **FACE** | **0.0 ± 0.2** | **-4.3 ± 4.6** | **-3.9 ± 5.2** | **-4.2 ± 5.2** | **0.4 ± 1.5** | **6.3 ± 3.2** |
|  | **proTELL** | **0.3 ± 0.9** | **-4.4 ± 5.2** | **-3.0 ± 5.9** | **-3.3 ± 6.0** | **1.2 ± 2.7** | **6.8 ± 3.4** |
|  | **SCI** | **0.5 ± 1.2** | **-6.7 ± 4.0** | **-6.5 ± 4.7** | **-6.8 ± 4.1** | **0.2 ± 1.1** | **7.4 ± 3.4** |
|  | **pooled** | **0.5 ± 0.8** | **-5.0 ± 2.9** | **-3.7 ± 3.1** | **-4.7 ± 3.1** | **0.6 ± 1.2** | **6.6 ± 2.5** |

Note: Pooled results refer to the average rating by pooling 4 associations within each type of associations (i.e., charity unfamiliar, CUF; charity familiar, CF; drug legalization group, D; gun/hunting rights advocacy group, GH). Bold texts indicate ratings for those associations finally selected for the pilot behavioral study and the current fMRI study. See the above section “vignettes used in the pilot rating task” for full names of all these associations.

**Table S5.**
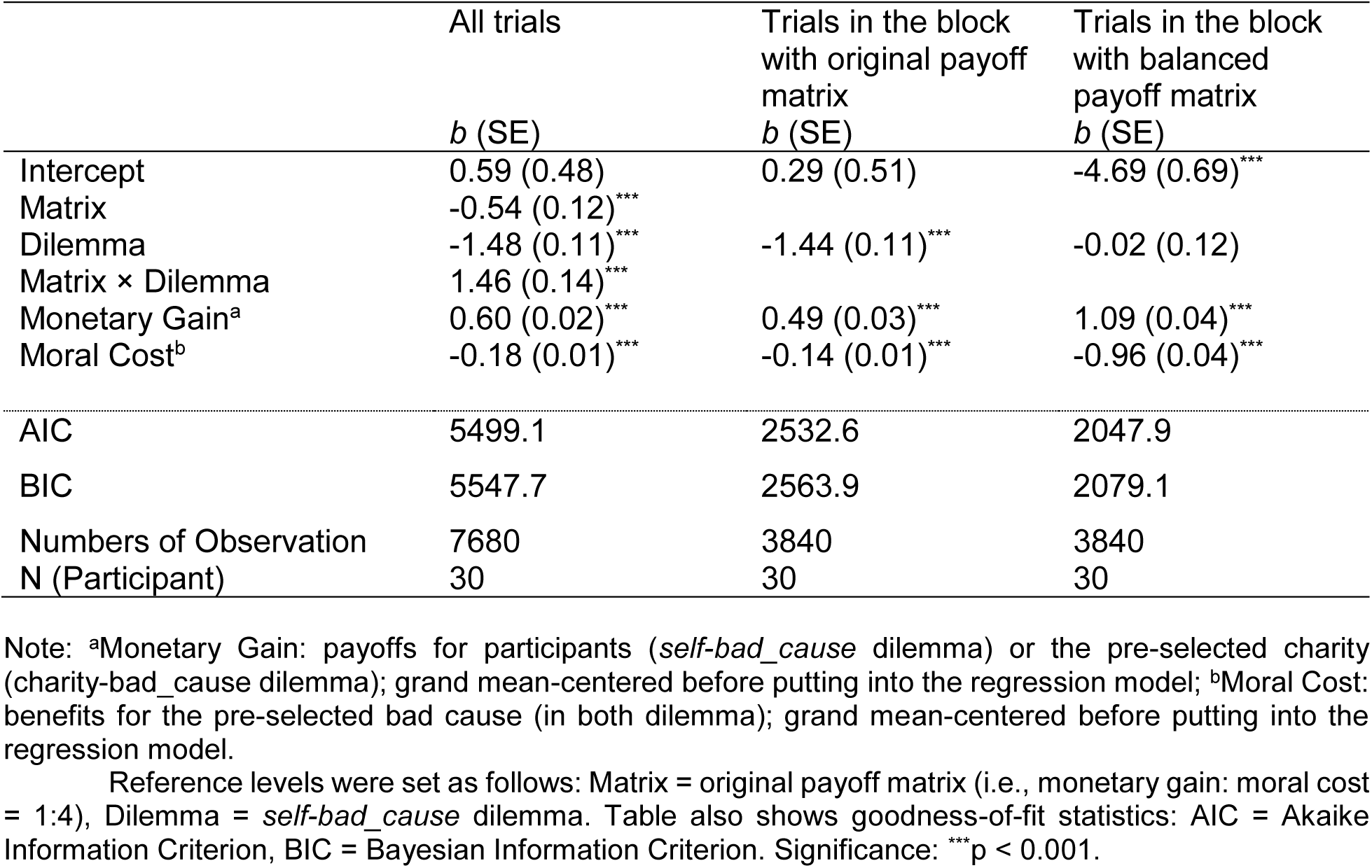
Output of fixed effects in mixed-effect logistic regressions predicting accept decisions in the pilot behavioral study

**Table S6.** Descriptive summary for the decision time (DT) and log-transformed DT (logDT; in ms) in the pilot behavioral study

|  |  | DT |  |  |  | logDT |  |  |  |
| --- | --- | --- | --- | --- | --- | --- | --- | --- | --- |
|  |  | Original |  | Balanced |  | Original |  | Balanced |  |
|  |  | self-bad_cause | charity-bad_cause | self-bad_cause | charity-bad_cause | self-bad_cause | charity-bad_cause | self-bad_cause | charity-bad_cause |
| Accept | Mean | 1143.6 | 1351.7 | 1067.8 | 1094.7 | 6.96 | 7.12 | 6.86 | 6.89 |
|  | (SD) | (355.7) | (350.6) | (400.0) | (363.8) | (0.33) | (0.32) | (0.36) | (0.34) |
|  | *N* | 28 | 28 | 28 | 30 | 28 | 28 | 28 | 30 |
| Reject | Mean | 1143.3 | 1134.4 | 1248.5 | 1268.7 | 6.95 | 6.95 | 7.05 | 7.06 |
|  | (SD) | (327.8) | (295.2) | (311.3) | (351.3) | (0.27) | (0.27) | (0.25) | (0.27) |
|  | *N* | 27 | 28 | 25 | 27 | 27 | 28 | 25 | 27 |

Note: we first calculated the individual-level mean DT and logDT in terms of specific decisions for each context of moral dilemma in each type of payoff matrix, then we calculated the group-level mean ( ± SD) based on the individual mean; Due to individual difference in decisions, the sample size (i.e., *N*) in original payoff matrix (i.e., monetary gain: moral cost = 1:4) and balanced payoff matrix (i.e., monetary gain: moral cost = 1:1) for each dilemma context of specific decisions is different.

**Table S7.** Output of fixed effects in mixed-effect linear regressions predicting logDT in the pilot behavioral study

|  | All trials | Accept trials | Accept trials<br>(original matrix) | Accept trials<br>(balanced<br>matrix) | Reject trials |
| --- | --- | --- | --- | --- | --- |
|  | *b* (SE) | *b* (SE) | *b* (SE) | *b* (SE) | *b* (SE) |
| Intercept | 6.87 (0.05)*** | 6.97 (0.06)*** | 6.95 (0.06)*** | 7.00 (0.07)*** | 7.01 (0.05)*** |
| Decision | -0.003 (0.02) |  |  |  |  |
| Matrix | -0.09 (0.02)*** | -0.09 (0.02)*** |  |  | 0.07 (0.02)*** |
| Dilemma | 0.16 (0.02)*** | 0.15 (0.02)*** | 0.15 (0.02)*** | 0.04 (0.01)** | 0.01 (0.01) |
| Decision × Matrix | 0.18 (0.02)*** |  |  |  |  |
| Decision × Dilemma | -0.10 (0.02)*** |  |  |  |  |
| Matrix × Dilemma | -0.12 (0.02)*** | -0.12 (0.02)*** |  |  | 0.02 (0.02) |
| Decision × Matrix ×<br>Dilemma | 0.10 (0.03)** |  |  |  |  |
| Monetary Gain<sup>a</sup> | 0.007 (0.002)*** | -0.02 (0.003)*** | -0.01 (0.005)** | -0.02 (0.003)*** | 0.02 (0.002)*** |
| Moral Cost<sup>b</sup> | -0.003 (0.001)*** | 0.001 (0.001) | 0.001 (0.001) | 0.02 (0.003)*** | -0.005 (0.001)*** |
| AIC | 5534.5 | 2658.8 | 929.2 | 1419.4 | 2169.5 |
| BIC | 5617.9 | 2708.5 | 960.2 | 1454.0 | 2219.8 |
| Numbers of<br>Observation | 7680 | 3676 | 1287 | 2389 | 4004 |
| N (Participant) | 30 | 30 | 29 | 30 | 29 |

Note: <sup>a</sup>Monetary Gain: payoff for participants (*self-bad\_cause* dilemma) or the pre-selected charity (*charity-bad\_cause* dilemma); grand mean-centered before putting into the regression model; <sup>b</sup>Moral Cost: benefits for the pre-selected bad cause (in both dilemmas); grand mean-centered before putting into the regression model.
Reference levels were set as follows: Decision = accept, Matrix = original payoff matrix (i.e., monetary gain: moral cost = 1:4), Dilemma = *self-bad\_cause* dilemma. Table also shows goodness-of-fit statistics: AIC = Akaike Information Criterion, BIC = Bayesian Information Criterion. Significance: \*p < 0.05, \*\*\*p < 0.001.

